# Model order reduction for left ventricular mechanics via congruency training

**DOI:** 10.1101/694075

**Authors:** Paolo Di Achille, Jaimit Parikh, Svyatoslav Khamzin, Olga Solovyova, James Kozloski, Viatcheslav Gurev

**Affiliations:** Healthcare and life sciences research, IBM TJ Watson research center, IBM research, Yorktown Heights, NY, USA; Ural Federal University, Yekaterinburg, Russia; Institute of Immunology and Physiology, Ural Branch of the Russian Academy of Sciences (UB RAS), Yekaterinburg, Russia

## Abstract

Computational models of the cardiovascular system and heart function are currently being investigated as analytic tools to assist medical practice and clinical trials. Recent technological advances allow for finite element models of heart ventricles and atria to be customized to medical images and to assimilate electrical and hemodynamic measurements. Optimizing model parameters to physiological data is, however, challenging due to the computational complexity of finite element models. Metaheuristic algorithms and other optimization strategies typically require sampling hundreds of points in the model parameter space before converging to optimal solutions. Similarly, resolving uncertainty of model outputs to input assumptions is difficult for finite element models due to their computational cost. In this paper, we present a novel, multifidelity strategy for model order reduction of 3-D finite element models of ventricular mechanics. Our approach is centered around well established findings on the similarity between contraction of an isolated muscle and the whole ventricle. Specifically, we demonstrate that simple linear transformations between sarcomere strain (tension) and ventricular volume (pressure) are sufficient to reproduce global pressure-volume outputs of 3-D finite element models even by a reduced model with just a single myocyte unit. We further develop a procedure for congruency training of a surrogate low-order model from multi-scale finite elements, and we construct an example of parameter optimization based on medical images. We discuss how the presented approach might be employed to process large datasets of medical images as well as databases of echocardiographic reports, paving the way towards application of heart mechanics models in the clinical practice.

## Introduction

Multi-scale finite element (FE) models of cardiac mechanics are being investigated as novel tools for analyzing medical imaging data and assisting personalized diagnostics in cardiac resynchronization therapy and disease monitoring [1–4]. Despite recent advances, personalizing FE simulations of heart mechanics to a specific clinical case still constitutes an open research problem. While most state-of-the-art models can capture the anatomical details of atria and ventricles from 3-D medical images (e.g., from CT or from the more costly MRI [5, 6]), significant computational resources are necessary to inversely estimate, when at all possible, the many unknown model parameters. Applying FE models in large scale analyses of clinical trials is, therefore, currently intractable. At the same time, it is widely accepted that a more mechanistic understanding of trial results could be greatly beneficial. For example, direct applications of models could provide illustrations of certain biophysical correlates of trial trends, thus aiding the interpretation of complex drug effects, such as intropes on cardiac function in heart failure patients [7–9].

Clinical evaluations often rely on echocardiography as their main source of cardiac functional and anatomical assessments, with analysts typically not having access to the raw acquisitions from the study, but rather relying only on succint clinical reports summarizing the measurements as a list of discrete indices. To more completely leverage information from echocardiography, multi-scale models would additionally exploit simple descriptions of anatomy provided by these limited ultrasound assessments. In such a framework, anatomical parameters would be estimated, and uncertainty quantified statistically by sampling (e.g., in a generalized polynomial chaos framework [10]). Again, due to complexity, the large computational expense connected to statistical approaches cannot currently be absorbed into the clinical application when FE models are employed directly. Instead in this study, we lay groundwork for a more feasible approach employing model order reduction, and a novel strategy to reduce 3-D FE models is the subject of this paper.

When using state-of-the-art biophysical models of heart, simulating even just a few beats can require hours to days of supercomputer time. Simplified models are therefore desirable, but they typically lack accuracy or interpretability. A clear example is provided by image-based models of the heart (see [11, 12] for two recent reviews), for which low-order analogues have typically relied on restrictive assumptions that fail to accurately represent the anatomy of ventricles and atria [13–16]. The lack of a clear bridge between imaged anatomies and model representations makes it difficult to link unknown parameters to radiological or physiological measurements. Thanks to basis simplifications introduced directly at runtime, advanced numerical techniques such as proper orthogonal decomposition [17–20], reduced basis functions [21, 22], and hyper-reduction [23] are able to speed up high-resolution computations by simplifying both the temporal and spatial solution spaces while at the same time maintaining a strong correspondence with their computationally expensive counterparts. These techniques provide significant advantages as they can rely on strong theoretical proofs of congruency to high resolution simulations, and also because they do not require design changes such as modifications to the base model’s parameterization. Obstacles to their adoption have been, however, their implementation complexity and the relatively contained speed-ups that such techniques can achieve. Although less rigorously, further speedups can be achieved by selecting *a priori* the subset of main global deformation modes assumed to drive contraction, and then by restricting the solution space accordingly [24].

Alternative strategies have proposed training purely statistical models (e.g., Gaussian Process regressions [25]) on a relatively limited number of high-resolution runs (e.g., see [26, 27] for applications in diastolic filling). Once fitted to a sampled training set, statistical models can be used as efficient surrogates for the computationally expensive simulations. These simulations can then be interrogated as needed by suitable optimization algorithms [28, 29]. Recent advances in co-kriging formulations [30] have also enabled to construct regressions that can leverage training on combined datasets comprising results from both low- and high-accuracy simulation results (e.g., see [31] for an application in cardiac electrophysiology). It is therefore apparent that while inferring model outputs from a statistical surrogate is computationally efficient, the accuracy of the approximations strongly depends on the quality of the training set, which might require a large computational cost to be built.

In this work, we demonstrate a method to circumvent some of the limitations of the above strategies by adopting a different hybrid approach. We introduce novel low-order (LO) models of ventricular mechanics that match global results from state-of-the-art FE simulations across multiple patients. The LO models comprise two classes of model components: first, “trained” black box transformations with parameters sized by machine learning to match model outputs to features of the detailed models, and therefore only indirectly linked to the underlying biophysics; and second, “physical” components, with parameters shared with the high-resolution biophysical model and therefore accessible to inference and prediction of underlying biophyiscal states, previously only available with the detailed models.

Fig 1 shows a schematic of the method. A typical FE model is composed of multiple elements, all of which in turn incorporate models of the biological units. Coupled interactions between FEs and the imposed boundary conditions ultimately result in the global outputs of the detailed model. The corresponding LO description accounts for only one or few biological units that are coupled to the global outputs through a series of transformations. In the simplest case, a transformation can be a linear operator (e.g., a scaling matrix or linear regression) defined by the set of “trained” parameters. Training is then performed by ensuring congruency between outputs of the LO and FE models upon a set of perturbed conditions.

**Fig 1.**
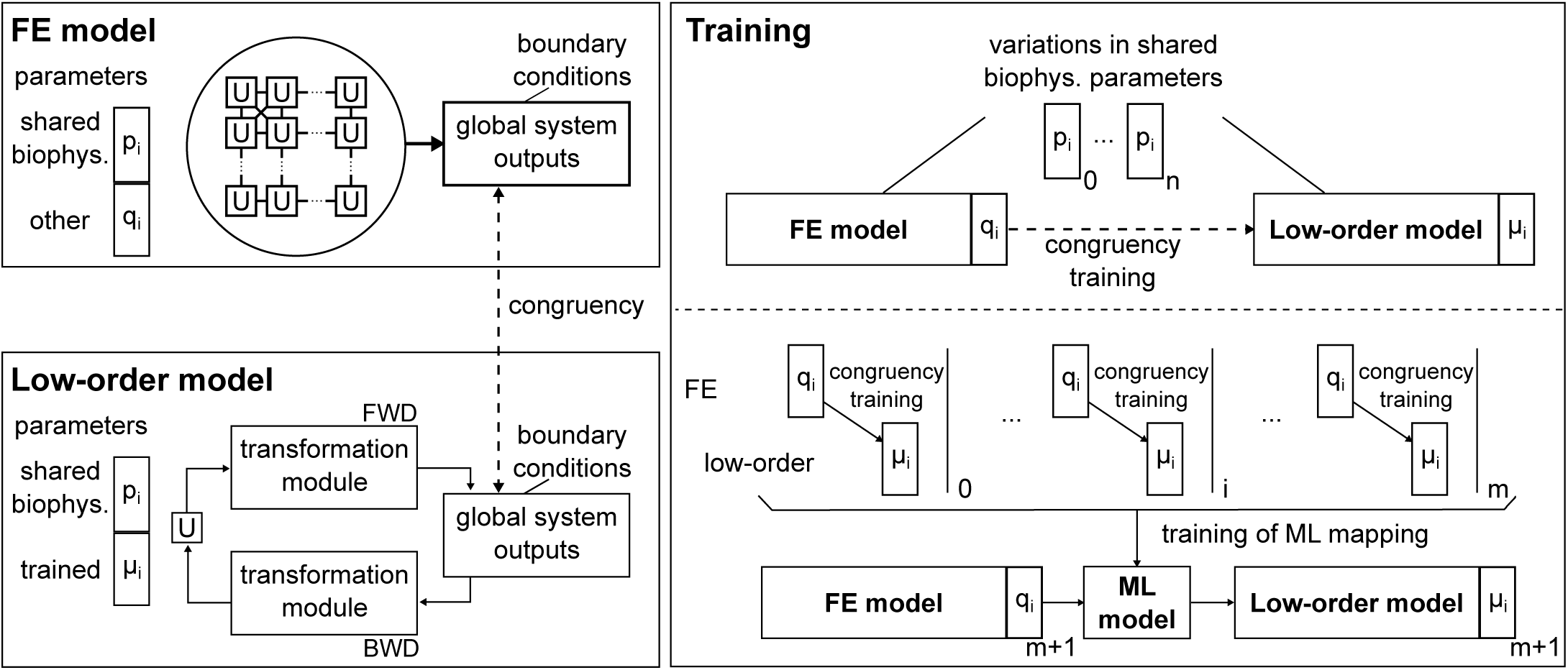
Schematic of low-order (LO) models and congruency training to FE simulations. The FE model on the top left accounts for the interconnections between cellular units (U) that cooperate to generate global outputs driven by “biophysical” parameters (i.e., *p*_*i*_ and *q*_*i*_). The LO model on the bottom left accounts instead for just a single U (modulated by the shared subset of biophysical parameters, *p*_*i*_) that is coupled to the global outputs via the transformation modules (modulated by the “trained” parameters, *µ*_*i*_). On the right, the training procedure is explained in more detail. Variations imposed on the “biophysical” parameters that are shared by both FE and LO models provide a set of perturbed simulations upon which the “trained” parameters are optimized to ensure congruency of global outputs. Once the procedure is repeated for a sufficient number of cases, a machine learning-based regression model can be used to infer “trained” parameters for additional cases.

As an example application, we construct LO models of 2 representative ventricles extracted from the publicly available Sunnybrook Cardiac MRI database [32]. After showing that the LO models are indeed congruent to their corresponding FE simulations upon varying training conditions, we test them under different simulated scenarios showing remarkable accuracy. Additional simulations performed on multi-unit LO models accounting for smoothly varying heterogeneities provide additional insights on how the LO models, despite their simplifications, can capture the global behaviors. We then explore the relationship between our “trained” phenomenological parameters and the anatomical features of the ventricles by training LO models for 106 “virtual” anatomies sampled from realistic distributions of geometric features. Finally, we employ the LO models as surrogates for the high-resolution FE simulations in an inverse optimization problem aimed at matching cardiac phase durations and ejection fraction measured from cardiac MRI.

## Materials and methods

### 3-D FE models of left ventricular mechanics

By integrating the cellular contraction model into a 3-D finite element formulation of soft tissue biomechanics, we sought to explore how the cooperative behavior of myocardial myocytes would translate at the macroscopic organ scale. FE models predict global outputs such as time-varying ventricular volume and pressure while also naturally accounting for the detailed anatomies of the heart and for the complex boundary conditions acting on it (e.g., the behavior of the downstream vasculature). In this work, multi-scale coupling between cellular and tissue models was implemented according to a stabilized mixed u-P formulation with ad hoc preconditioning and adapted for P1/P1 elements [27, 33]. Briefly put, coupling between cellular and elemental stresses and strains occurred implicitly at the Gauss point level. Passive contributions from extracellular matrix and from myocyte structural proteins were described as a hyperelastic behavior following the constitutive model by Usyk et al. [34] (see Supplemental Material for more details). To simulate also active contraction, calcium transients triggering myocyte response had to be synchronized to the pacing dictated by heart electrophysiology. Rather than simulating in detail the generation and propagation of action potentials, we followed a simplified decoupled approach where information on arrival timing of the electric impulse wavefront was computed by solving the Eikonal equation with anisotropic conductivities [35]. Specifically, myocardial fibers were assumed to have 2 times higher conductivity than myocardial sheets and 4 times higher conductivity than mutually normal cross-fibers. Studies on mapping electrical activation of the heart suggest that Purkinjie fibers might resurface at a small number of endocardial foci to initiate impulse propagation [36]. We selected, therefore, 4 small groups of endocardial elements as zero-time regions in the eikonal solution. In polar coordinates 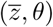, the set of 4 prescribed activation foci was 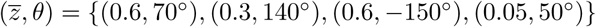, where the first value of each tuple indicates whether the activation focus was closer to the base (i.e., 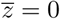) or to the apex (i.e., 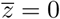), while the second value represented the circumferential coordinate (in degrees) measured counterclockwise around the longitudinal axis of the ventricle starting from the approximate center of the left ventricular free wall (*θ* = 0°).

### Computational domains and boundary conditions

While the FE formulation could be in principle applied to arbitrarily complex geometric domains such as detailed multi-chamber models, this study targeted only the behavior of left ventricles (LVs). Specifically, we employed a parameter axisymmetric representation of ventricular anatomy that was recently employed to automatically process the Sunnybrook Cardiac MRI database [27, 32]. Figure 2 shows axial and longitudinal cross-section profiles of the two LV geometries that were selected as representative of the normal (N) and heart failure (HF) subject groups included in the database. To at least partially correct for the fact that all imaged ventricle configurations are subjected to non-negligible loads (e.g., due to intraventricular pressure and external boundary conditions), we first “unloaded” the geometries reconstructed from MRI using Gaussian Process regression under the assumption of a 10% mid-wall end-diastolic strain for the N geometry and a 15% mid-wall strain for the HF one (see leftmost and central columns for cross-section profiles at end-diastole and after unloading, respectively) [27]. The 2 idealized anatomies were then discretized into 3-D meshes of 34 590 (N) and 49 121 (HF) linear tetrahedral elements (see righmost column). Ventricle bases (marked in yellow) were prevented to move out of plane, while a 5 mm-wide stripe of epicardial elements (marked in red) closest to base was fully constrained to prevent rigid motions. Pressure and volume couplings between the left ventricle, the left atrium and the distal vasculature was weakly enforced at the endocardial surface as a Neumann boundary condition (marked in blue).

**Fig 2.**
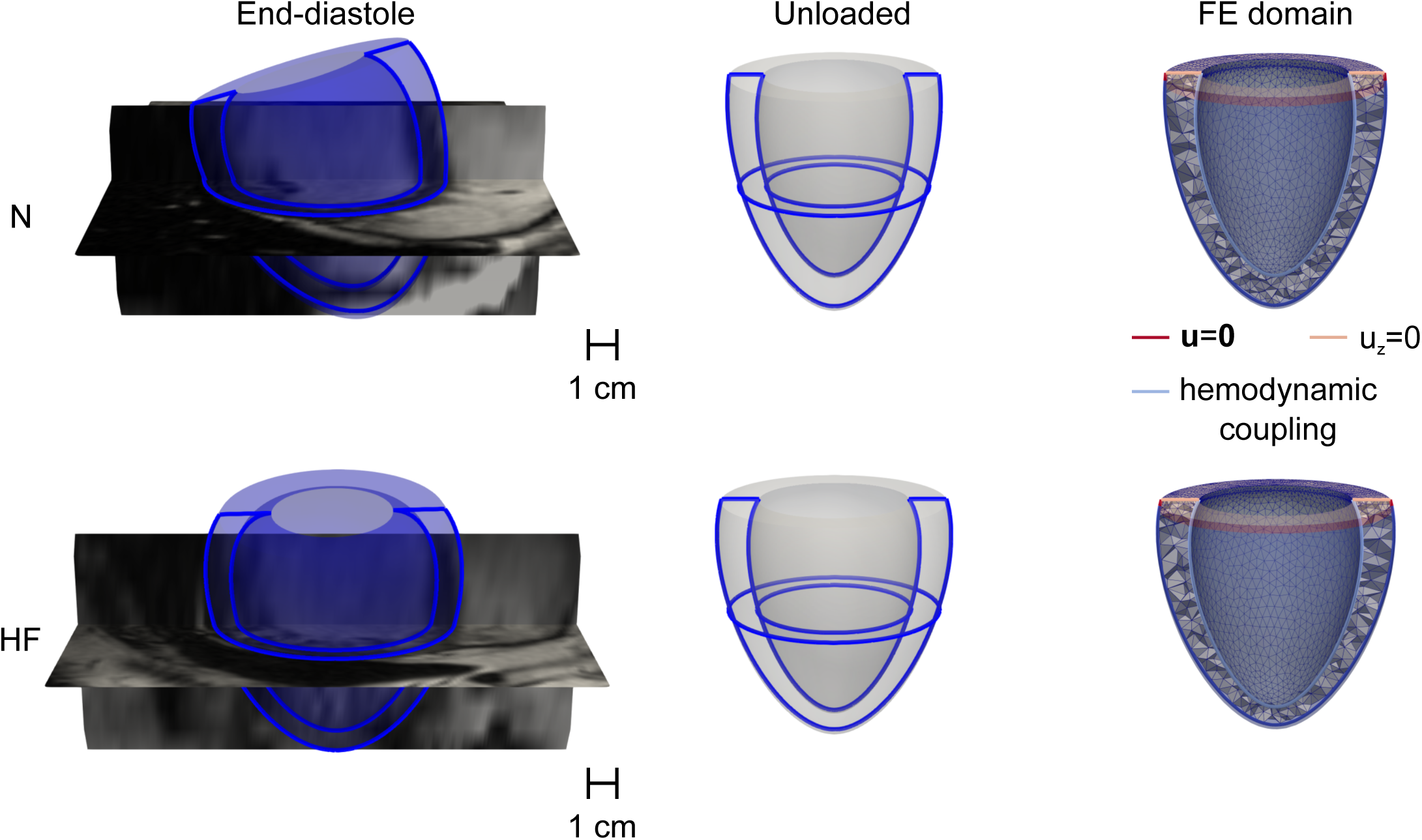
Discretization of the two computational domains considered as representative of a normal (N) and heart failure (HF) LVs extracted from the Sunnybrook Cardiac MRI database. Leftmost column: cross-sections profiles matching end-diastolic configurations are overlayed with corresponding MRI slices (see Table 1). Central column: cross-section after unloading via GP-regression under the assumption of 10% (15%) mid-wall stretch for the N (HF) geometry. Rightmost column: discretization of the unloaded geometries into finite element meshes. Overlayed colored segments mark the different boundary conditions applied: zero displacement within a 5-mm wide stripe of epicardial elements adjacent to base (red), zero axial displacement at base (yellow), and weakly imposed hemodynamic pressure-volume coupling acting on the inner surface of the ventricle (light blue).

### Low-order models of ventricular mechanics

Parallel schematics of the high-resolution FE models and the here proposed corresponding LO counterparts are reported in Fig 3. In terms of formulation, the FE models relied on a PDE-based description of mechanical equilibrium coupled via hemodynamic boundary conditions to a simplified representation of atrial pressure (*P*_*at*_) and to a 3-element Windkessel model of the downstream vasculature with additional resistance to split aortic valve *R*_1_, aorta *R*_2_, and peripheral artery resistance *R*_3_. In training the LO model, we used a simpler configuration with *R*_2_ = 0, combining aortic and peripheral system. In the model, the capacitance of the peripheral system is denoted as *C*. The LO models maintained identical hemodynamic components, but also simplified the description of the computational domain (Ω) to a set of algebraic relationships linking cellular active tension (*T*_*a*_) to left ventricular pressure (P), and cellular elongation (*λ*) to left ventricular volume (V).

**Table 1.** 6-parameter axisymmetric description of ventricular geometries at end-diastole and after virtual unloading. In the rightmost column, left ventricular volumes at each configuration.

| Group | configuration | $R_b$<br>(mm) | $L$<br>(mm) | $Z$<br>(mm) | $H$<br>(mm) | $e$ | $\Psi_0$<br>(deg) |
| --- | --- | --- | --- | --- | --- | --- | --- |
| N | end-diastole | 38 | 8.2 | 52 | 11 | 0.76 | -61 |
|  | unloaded | 35 | 9.6 | 54 | 12 | 0.70 | -53 |
| HF | end-diastole | 42 | 8.2 | 49 | 6.0 | 0.71 | -78 |
|  | unloaded | 38 | 10 | 49 | 7.7 | 0.66 | -64 |

**Fig 3.**
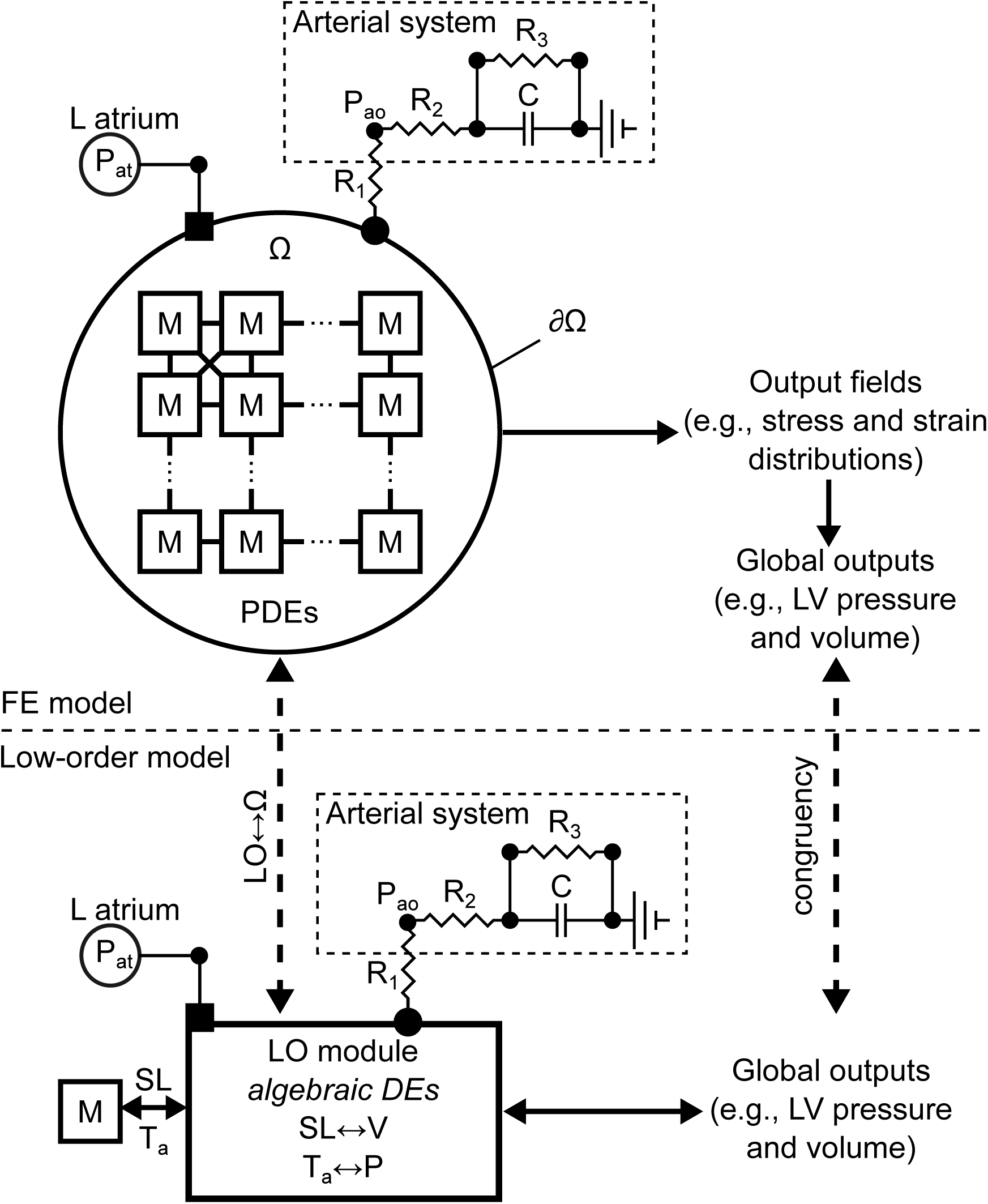
Parallel between between schematics of finite element and novel LO models of ventricular mechanics. Note how the architecture and structural organization of the many myocyte units (M) of the FE models is described in the LO framework by a transformation module coupled to a representative myocyte and to lumped parameter models of atrium and cardiovasculature. Parameters for the LO module are sized via congruency training to ensure good correspondence between global outputs (e.g., LV pressure and volume traces) among the FE and LO descriptions.

#### Passive behavior

The hyperelastic passive mechanical behavior of ventricles was modeled by the 3-D strain energy function presented in [34] (see Supplemental Material for more details). To reproduce an equivalent behavior, our LO models were endowed with empirical relationships mimicking the combined effects of nonlinear hyperelasticity and ventricular geometry. Specifically, we considered separate formulations for the responses observed at volumes higher and lower than the unloaded volume *V*_0_,

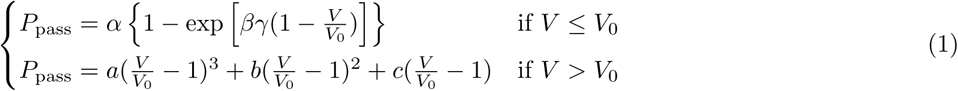

where *α, β, γ* are coefficients modulating the pressure-volume exponential behavior upon compression; *a, b, c* are coefficients of the 3^rd^-order polynomial describing the ventricle under passive tension. The differential treatment of the behavior under compression or tension was necessary to ensure good fits of the LO model behavior to the FE simulations.

#### Active behavior

Coupling between active myocyte response and time-varying ventricular hemodynamics is a pivotal component of the LO models introduced here. The relationships driving cell action can be summarized as

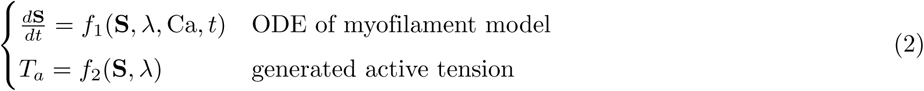

where **S** is a vector containing the time-varying state variables of an ODE model of myofilament contraction that depends on calcium concentration, Ca, and on strain of the myofilament, *λ*; state variables and strain also uniquely determine the active tension generated by the myofilament *T*_*a*_ (see Supplemental Material for the specific *f*_1_ and *f*_2_ used for this work).

The relationships in (2) need to be opportunely transformed to incorporate the role played by the coordinated contraction of the left ventricle, with correct sizing of the transformations being of great importance for building LO models for a given ventricular anatomy of interest. The relationships linking cellular strain to ventricular volume and active tension to ventricular pressure are given by

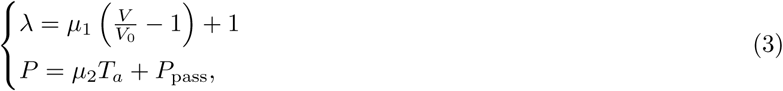

where *µ*_1_ and *µ*_2_ are opportune scaling coefficients of the active component of the low order model, and *P*_pass_ follows what shown in (1).

#### Congruency training

We now describe the strategy used to find the overall best-fit coefficients for the linear transformations in (3) from high-resolution FE runs. To ensure that the LO models maintained correspondence to their FE counterparts over a wide range of myocyte lengths and downstream hemodynamic resistances, we considered a set of 3 active and 1 passive training simulations encompassing varying hemodynamic conditions. More specifically, to extract the passive coefficients {*α, β, a, b, c*} we fitted the LO model predictions to a FE pressure-inflation test performed using an intraventricular pressure range of 0-4 kPa. Note the estimation *α*, and *β* was done under the assumption of neutral role of *γ* (i.e., *γ* = 1), as we defined *γ* to capture differential behavior upon compression conditions (see (1)), which are not tested upon passive inflation. Best-fit values of the remaining parameters {μ_1_,μ_2_ ,*γ*} were instead found from active simulations accounting for different combinations of atrial pressure and Windkessel parameters (see Table 2). In other words, for any ventricular geometry defined by {*R*_*b*_, *L, Z, H*, Ψ_0_, *e*}, we solved two sequential problems to maximize correspondence between global outcomes of the FE and the LO models. A first minimization problem leveraged the results of the passive inflation simulation,

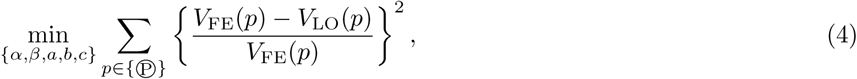

where the subscripts _FE_ and _LO_ are used to label the left ventricular volumes predicted by the FE and LO models, and *p* is a set of left ventricular pressures considered in the passive simulation ℗. The optimal best-fit coefficients for the passive material behavior were then used in a second minimization problem seeking the active “trained” parameters,

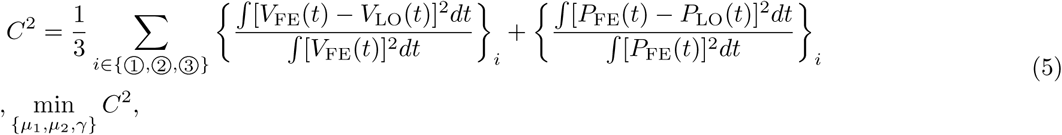

where the discrepancies between global outputs *P* and *V* are computed for all of the active training simulations (i.e.,{➀,➁,➂}, see Table 2). To solve the problem in (4) we used a standard curve fitting procedure based on the Levenberg-Marquardt algorithm available in SciPy. Solving (5) was less straightforward, as it required iterative runs of the LO model to compare global outputs during active contraction. To exploit parallel execution on multicore machines, we used a custom implementation of the OrthoMADS algorithm [37], with search steps interrogating an iteratively updated kriging surrogate to speed up convergence [28, 29].

**Table 2.** Boundary conditions for training simulations

| Simulation | $P_{at}$<br>(kPa) | $R_1$<br>(kPa ml<sup>-1</sup> s) | $R_2$<br>(kPa ml<sup>-1</sup> s) | $R_3$<br>(kPa ml<sup>-1</sup> s) | $C$<br>(kPa<sup>-1</sup> ml) |
| --- | --- | --- | --- | --- | --- |
| Ⓟ | 0-4.0 | — | — | — | — |
| ① | 0.33 | 0.0015 | — | 0.20 | 8.0 |
| ② | 0.66 | 0.0015 | — | 0.05 | 8.0 |
| ③ | 1.5 | 0.0015 | — | 0.20 | 8.0 |

### Multi-element low-order models

In a simulated heartbeat, the many cell units of a high-resolution model undergo deformations that may vary based on their anatomical location (e.g., across the cardiac wall or along the apex-base direction). As a result, even for the same computational domain, cellular models of active contraction may operate at heterogeneous sarcomere lengths that cannot be explicitly accounted for a single-cell LO description. To incorporate heterogeneity and evaluate approximation errors, we then extended the LO models to account for multiple interconnected elements that share the same left atrium and Windkessel models. Figure 4 shows schematics of the two considered multi-element configurations, with LO submodules connected either in parallel (see panel A), or in series (see panel B). Outputs from each configuration were then compared to a single-element LO model that was adapted via congruency training to maximize correspondence between global outputs of the single-element and multi-element models. Congruency training was carried out solving (5) and the error analysis was repeated for 3 different atrial pressures, *P*_*at*_ = 0.3, 0.6, and 1 kPa.

**Fig 4.**
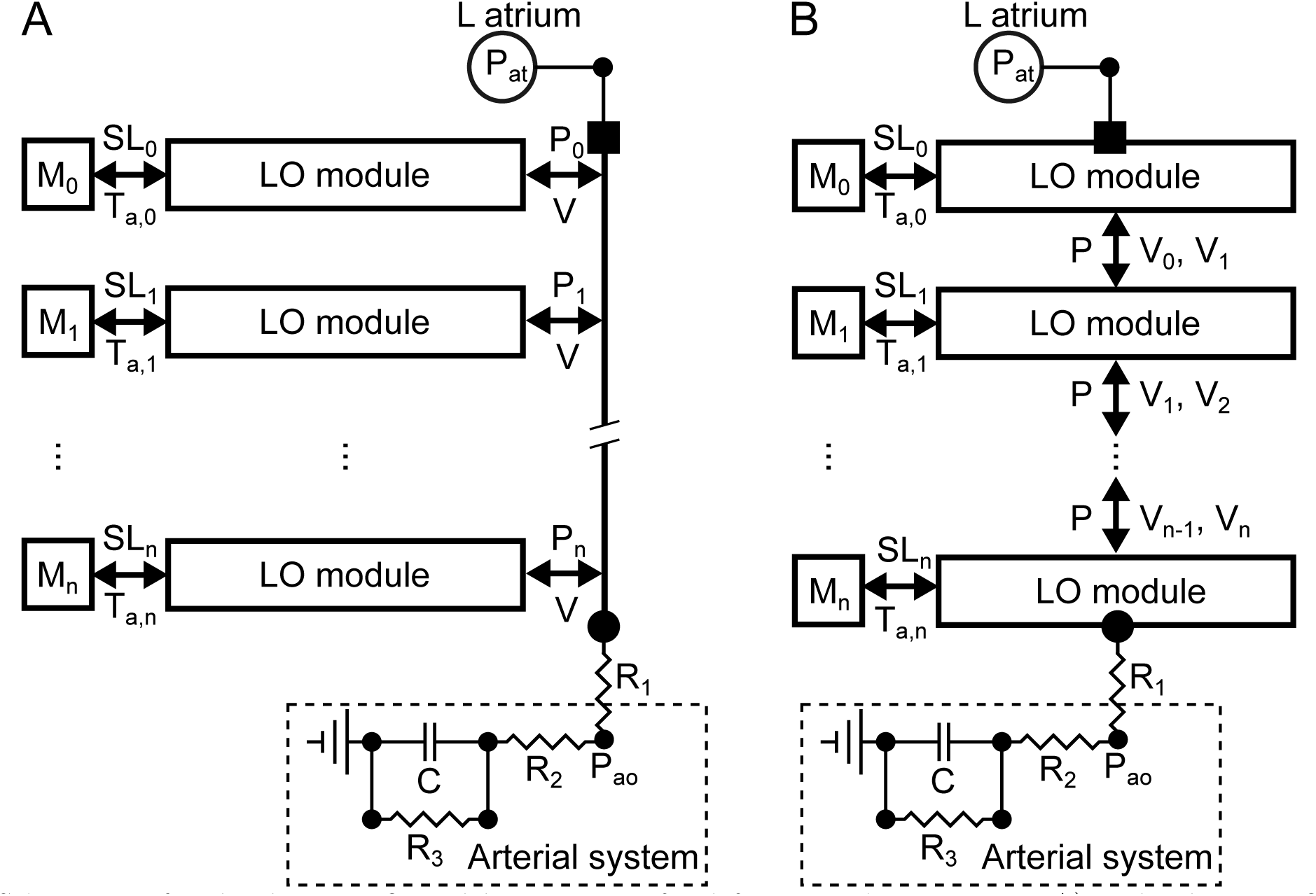
Schematics of multi-element LO models accounting for deformation heterogeneity. A) Multi-element LO models with subcomponents connected in parallel share the same ventricular volume, *V*, but operate at different sarcomere lengths (i.e., generate different active tension, *T*_*a,i*_) due to varying *µ*_1_ values. B) Multi-element LO models with components connected in series operate under the same ventricular pressure, *P*, but they each contribute a different volume subportion, *V*_*i*_, to the total volume, *V*. Volume magnitude and general behavior of each series element is modulated by weighing the LO parameters {*α, a, b, c, µ*_2_}.

#### Parallel configuration

The parallel model can be viewed as the combination of LO subcomponents that share the same ventricular volume *V* but operate at different sarcomere lengths because of non-uniform *µ*_1_ values. In this case, equation (3) takes the form

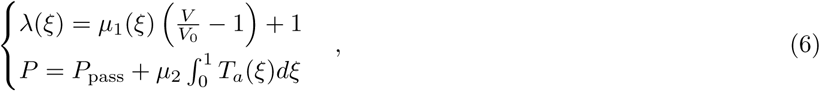

where *λ* and *T*_*a*_ are the stretch ratio and the active tension of the LO submodels, respectively, and *ξ* ∈ [0, 1] indicates the discretization of the system into subcomponents. To include heterogeneity, the *µ*_1_ parameter was imposed to vary linearly within the [*µ*_1_(*ξ* = 0), *µ*_1_(*ξ* = 1)] = [0.01, 0.179] range. We overall considered 256 LO subcomponents.

#### Series configuration

The series model combines LO subcomponents each contributing a volume portion, *V* (*ξ*), to the global left ventricular volume, *V*, while operating under the same ventricular pressure, *P*. The weights *w*(*ξ*) introduce heterogeneity in the series multi-element model by modulating the effects of a subset of LO parameters, {*α, a, b, c, µ*_2_}. Accordingly, equation (3) takes the form

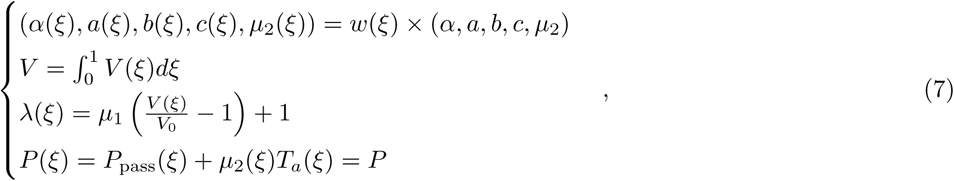

where *ξ* ∈ [0, 1] indicates the discretization of the system into subcomponents. The *w*(*ξ*) weights were imposed to vary linearly within the range [*w*(0), *w*(1)] = [0.6, 1.0]. Overall, the model was discretized into 256 LO subcomponents connected in series.

### Regression for congruency coefficients

While the minimization problems (4) and (5) were solved for any given geometry on the basis of 4 high resolution runs, the computational burden would scale up quickly in situations where many anatomies might be of interest. For this reason, we built a Gaussian Process regression model mapping the geometric parameters of the left ventricle (i.e., {*R*_*b*_, *L, Z, H*, Ψ_0_, *e*}) to the 8 parameters of the LO module (i.e., {*α, β, γ, a, b, c, µ*_1_, *µ*_2_}). To train the regression model, we randomly sampled 150 virtual anatomies uniformly distributed over the range of geometric features observed for the patients included in the Sunnybrook Cardiac MRI datasets [27]. 3-D high-resolution simulations were run for all geometries and under all training conditions (see Table 2). To ensure meaningful fits also for the *γ* parameter, which acts only when LVV is smaller than *V*_0_, we excluded from subsequent training the cases for which the left ventricle did not contract enough to reach 85% of the unloaded volume *V*_0_ under hemodynamic conditions ➁, which imposed a minimal afterload constraint. Efficacy of the Gaussian Process regression model in learning the behavior of the LO indices was evaluated via 5-fold cross-validation.

### Inverse problem solution: fit to MRI data

To test the efficacy of our LO models as convenient surrogates of high-resolution 3-D FE counterparts, we employed the overall model reduction framework to solve an inverse parameter identification problem. More specifically, we sought values for the model parameters that would ensure an optimal match with the ejection fraction and cardiac phase durations measured from the Sunnybrook Cardiac MRI scans [27, 32]. Knowing that heart failure might likely affect both calcium handling and overall contractility, we chose to optimize the *τ*_2_ and *S*_*a*_ parameters, which modulate the duration of the calcium transient in the cellular unit and the overall efficiency of contraction, respectively. At the same time, we also penalized deviations of central aortic blood pressure from the normal range 85-110 mmHg by adapting the hemodynamic parameters of the Windkessel submodel, i.e., the aortic resistance *R*_2_, the distal resistance *R*_3_, and the capacitance *C*. We used the same value for the aortic valve resistance as during the congruency training simulation (i.e., *R*_1_=0.0015 kPa ml^−1^ s). Finally, we assumed that the left atrium would contribute to diastolic relaxation by providing a progressively decreasing blood pressure. For this, we employed an electrical circuit analogy and represented the atrium as a capacitor. The capacitance value, *C*_*at*_, was considered unknown and optimized together with {*τ*_2_, *S*_*a*_, *R*_2_, *R*_3_, *C*}.

As in 5, the inverse optimization problem was solved via OrthoMADS, leveraging parallel execution. Comparison analyses were finally carried out with respect to the pressure and volume traces computed for the optimal set of parameters.

## Results

In our approach, the simulated global behavior is reproduced at a reduced computational cost by transforming the outputs of a modeled cellular subcomponent. To determine a suitable functional form for the cell-to-global transformations, our first analyses investigated the relationships between local and global biomechanical measures in active FE simulations. Fig 5 shows LVV plotted against mid-wall stretch in the fiber direction (*λ*_mw_) in simulations run under the same boundary conditions used for congruency training (i.e., ➀, ➁ and ➂in Table 2). Probing locations were taken across the ventricular wall at half of the apex-to-base distance and averaged over 40% and 60% of thickness. As expected, simulated perturbed conditions led to pronounced changes in local stretch for both the N (see black lines) and the HF (see gray lines) cases, with stretch ranges within 0.88-1.21. A least-square error fit revealed that the relationship between LVV and *λ*_mw_ could be reasonably assumed to be linear for both considered geometries (R^2^=0.99).

**Fig 5.**
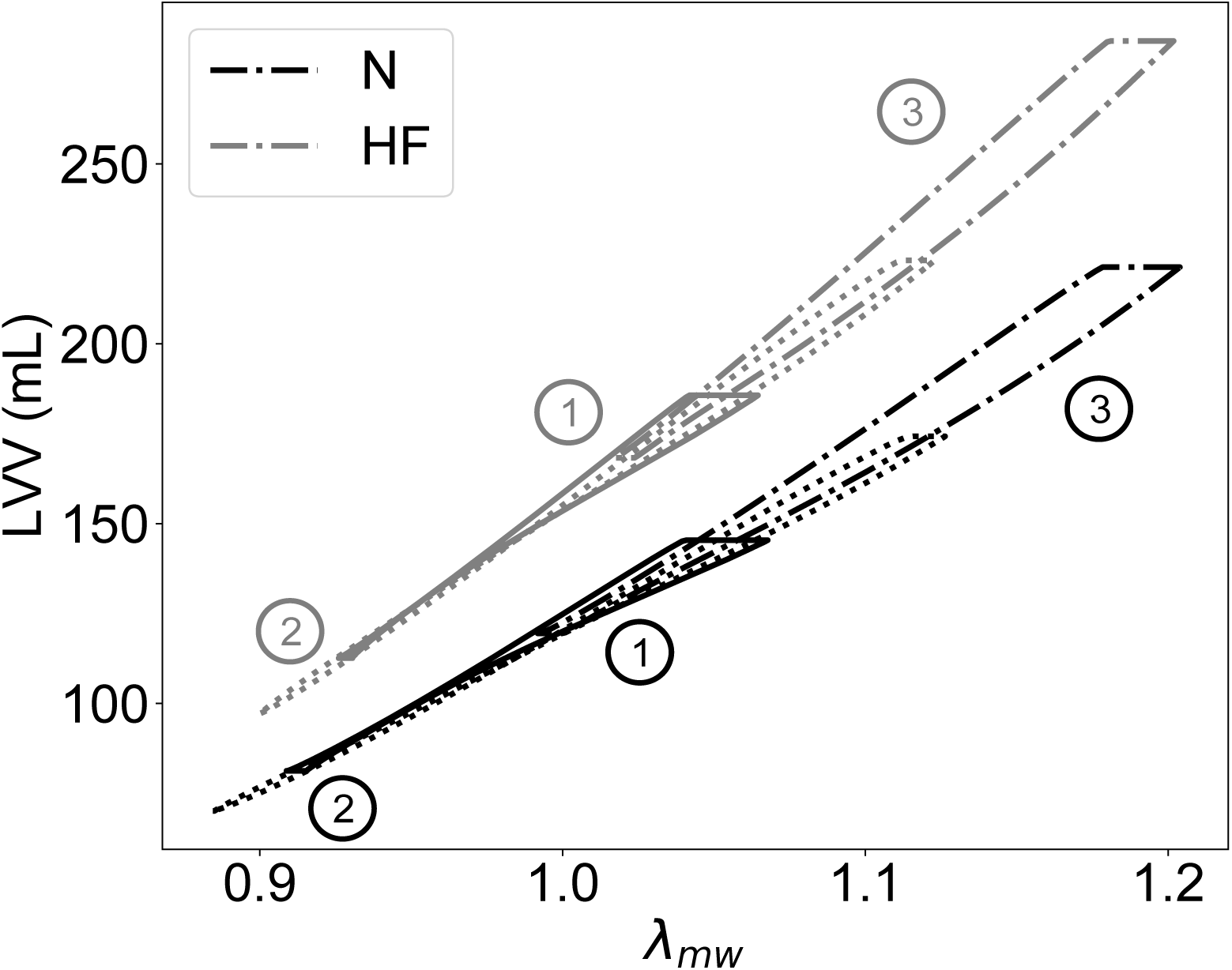
Quasi-linear relationships between left ventricular volume, LVV, and mid-wall stretch, *λ*_mw_ in the fiber direction for both the N and HF geometries. The approximately linear relationship (R^2^=0.99) between LVV and *λ*_mw_ was preserved over a wide range of perturbed boundary conditions (i.e., ➀, ➁ and ➂ see Table 2 and text for more details).

This relationship between simulation outputs at the global (LVV) and local (*λ*_mw_) levels caused us to adopt simple linear forms for the transformations in (3). Fig 6 shows the outputs from high-resolution FE (solid lines) overlayed with outputs from LO models (dashed lines) after parameter optimization via congruency training. Best-fit values for the trained parameters {*µ*_1_, *µ*_2_, *γ, α, β, a, b, c*} are reported in Table 3 for the N and HF ventricular geometries.

**Fig 6.**
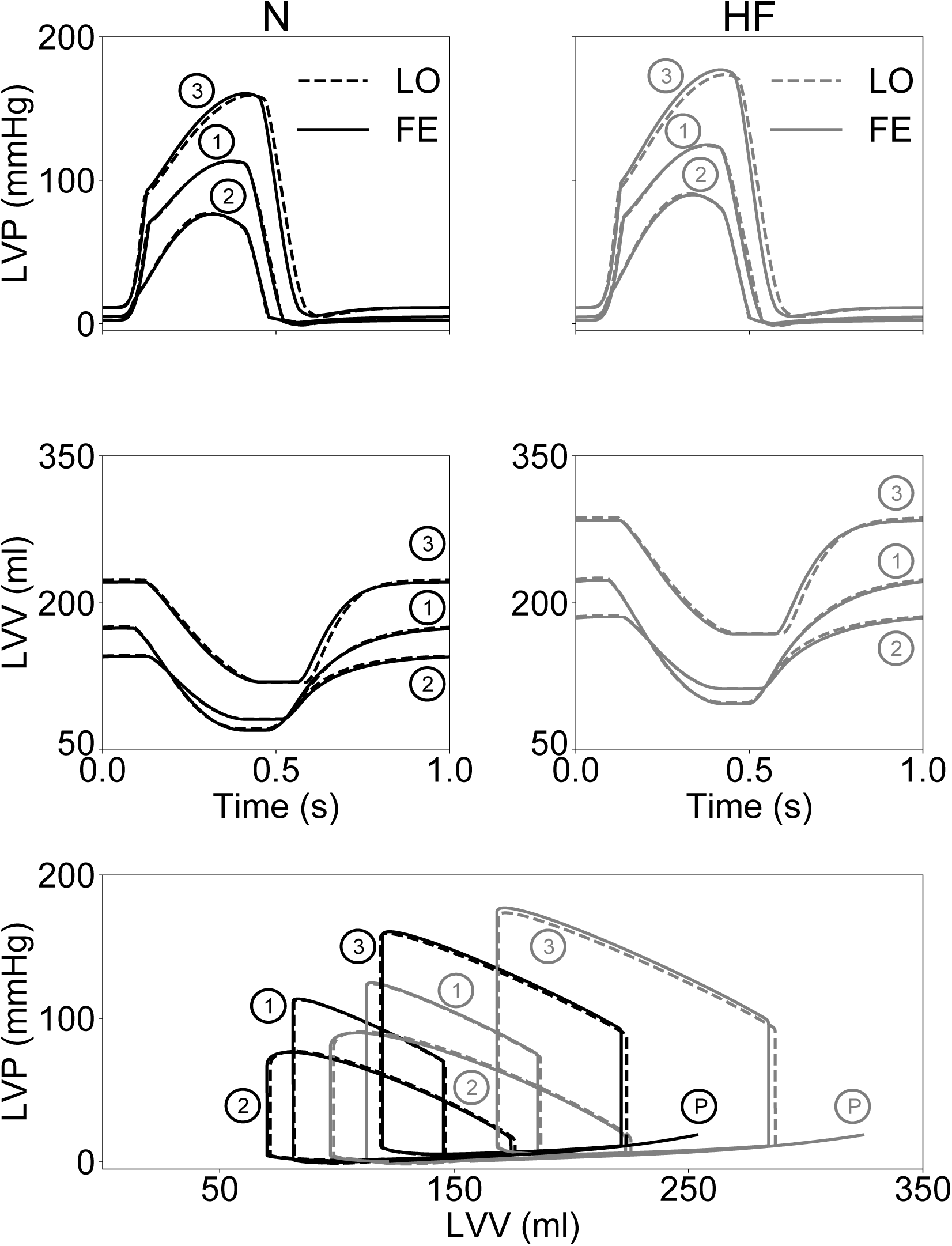
Correspondence between finite element (FE) and low-order (LO) models after congruency training of passive and active behaviors. Top row, left ventricular pressure (LVP) traces over the course of a cardiac cycle show an overall good match between active simulations of the two classes of models for both considered anatomies. Central row, similar to top row but for left ventricular volume traces over the course of a cardiac cycle, showing almost complete overlap in all cases. Bottom row, same traces of the rows are plotted as PV loops overlapped to the passive inflation tests ℗, showing how the LO model recapitulates well also passive behavior over an extended ranges of pressures and volumes.

**Table 3.** Best-fit parameter values from congruency training for the two geometries considered. *µ*_1_, *µ*_2_ and *γ* parameters were estimated via active training simulations (➀, ➁ and ➂) once the remaining parameters were constrained from the passive inflation test. ℗

| Geometry | Active training |  |  | Passive training |  |  |  |  |
| --- | --- | --- | --- | --- | --- | --- | --- | --- |
| | $\mu_1$ | $\mu_2$<br>(kPa) | $\gamma$ | $\alpha$<br>(kPa) | $\beta$ | $a$<br>(kPa) | $b$<br>(kPa) | $c$<br>(kPa) |
| N | 0.191 | 0.183 | 4.97 | 0.640 | 1.39 | 1.70 | -1.41 | 1.64 |
| HF | 0.190 | 0.173 | 5.17 | 0.571 | 1.47 | 1.77 | -1.47 | 1.61 |

The perturbed hemodynamic parameters considered during training affected significantly global performance both in terms of ejection fraction (observed ranges within 44%-60% and 40%-60% for the N and HF geometries, respectively), and peak systolic LVP (observed ranges within 77-158 mmHg and within 91-174 mmHg for the N and HF geometries, respectively). Note that LO and FE models employed the same values for all the “physical” parameters modulating myocyte contraction and hemodynamic coupling. Outputs from the two classes of models matched closely, with small discrepancies appreciable between pressure traces (see top row of Fig 6) and almost complete overlap occurring at the volume level (see central row in Fig 6). The largest, but still contained, mismatch was appreciable for simulation ➂ in the HF case (see gray lines in the top right panel Fig 6). For both classes of models, simulated pressure-volume loops are shown in the bottom row overlayed to results from the passive inflation tests run to optimize the LO coefficients {*α, β, a, b, c*}. Both relations in (1) resulted in good matches during passive filling, but the polynomial form provided better accuracy at lower pressures.

Despite supporting our choice for the functional forms (3), the results of Fig 6 did not provide any indications on whether the LO models would show congruent behavior also in scenarios not directly considered during training. To assess performance under more general conditions, we then compared global outputs of the two classes of models when independently varying sets of parameters modulating cellular action. Meanwhile, the trained parameters of the LO models were maintained fixed to the optimal values obtained via congruency training without further tuning (see Table 3). Fig 7 compares LVP (first and third rows) and LVV traces (second and fourth rows) obtained in the additional simulated scenarios. The first column shows effects of varying the maximum velocity of muscle shortening, *V*_*max*_, by simultaneously modulating the attachment and detachment rates of crossbridges, i.e., *f*_*XB*_ and *g*_*XB*_, respectively. The 3 shown simulations account for 50% rate variations above and below the baseline values used for training. Altering crossbridge dynamics resulted either in faster (for increased *f*_*XB*_ and *g*_*XB*_) or slower (for decreased *f*_*XB*_ and *g*_*XB*_) contractions during systole, as appreciable from the different LVV trace slopes observed during ejection. The LO results (see dashed lines) followed closely their FE counterparts (see solid lines) for both the N (top two rows) and HF (bottom two rows) cases, and with mismatches limited to the isovolumic relaxation phase, when ventricles operated under compression behavior.

**Fig 7.**
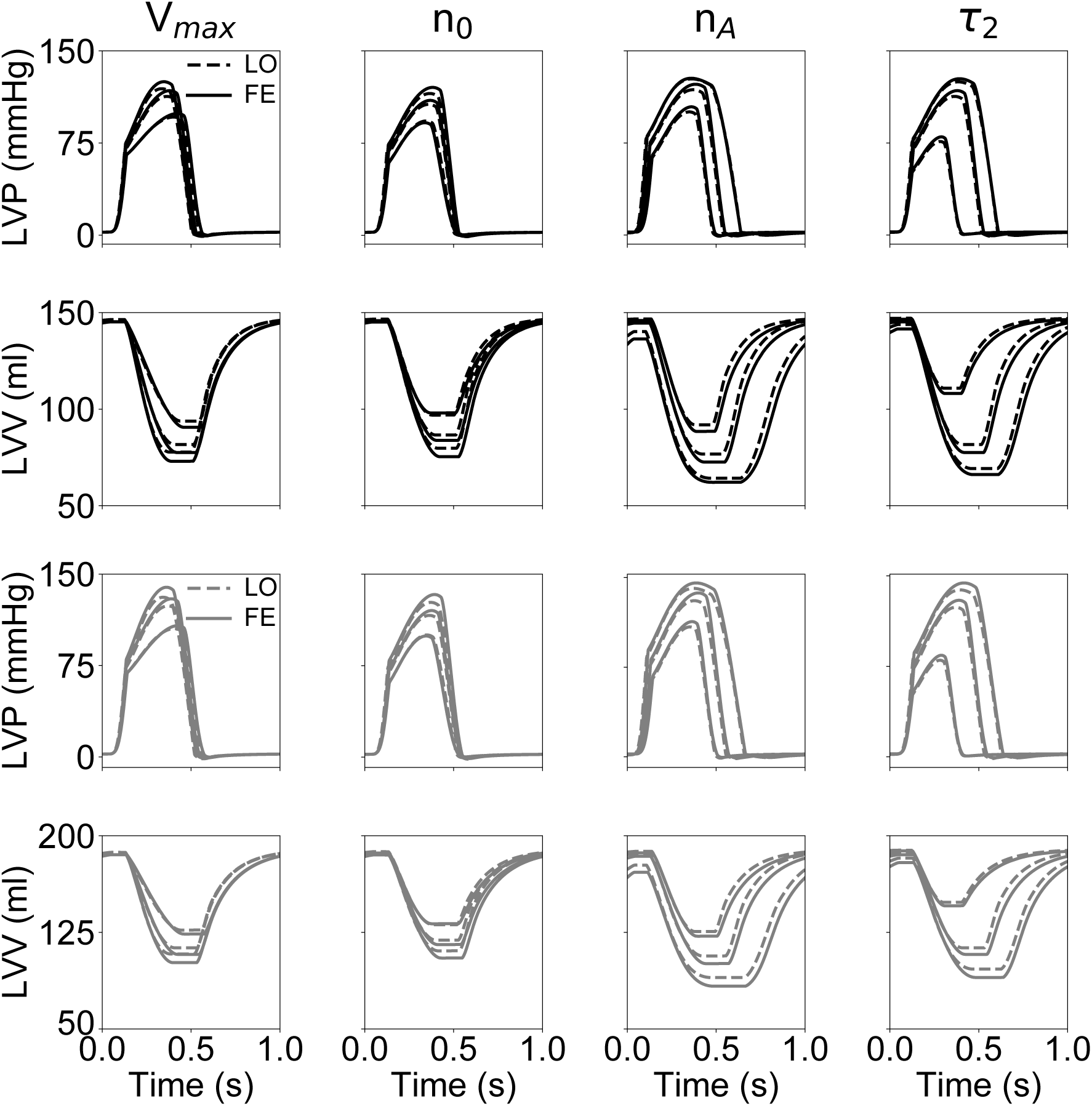
Reduced order models maintain good congruency to 3-D FE models even for cases not included in the training set. Top 2 rows, pressure and volume traces simulated via FE (solid lines) and LO (dashed lines) over the course of a cardiac cycle for the normal geometry N. Plots in each column show simulated behavior resulting from changes in one of the following parameters: *V*_*max*_, maximum velocity of contraction; *n*_0_, regulating the slope of the length-dependence function for both thick and thin filaments; *n*_*A*_, Hill coefficient of curve governing myofilament cooperativity; *τ*_2_, characteristic time of calcium transient decrease (see text and Supplemental Material for more details). Two bottom rows, same as above but traces were simulated by considering the HF geometry (see gray lines).

For a second scenario, we compared LO and FE simulation results for varying length-tension relationships. When progressively increasing *n*_0_, i.e., the minimum stretch required for myocytes to generate active force, ventricles generated progressively less force and contracted less. Maximum relative mismatch (i.e., absolute pointwise difference normalized by pointwise mean value) in LVP and LVV traces between FE and LO was ~ 3% on average. The third scenario is again depicted in Fig 7 (see third column) and targeted the *n*_*A*_ parameter, which defines coperativity in the interaction among fraction of troponin with bound calcium, tropomyosin state, and crossbridge dynamics (see Supplemental Material for more details). Increasing *n*_*A*_ resulted in smaller forces of contraction (i.e., larger end-systolic volumes) combined to faster ejection and isovolumic relaxation phases. These findings can be attributed to crossbridges being recruited at lower calcium levels when *n*_*A*_ is low, while increasing *n*_*A*_ resulted in crossbridges more quickly reaching their maximum generated force. Even for this set of simulations, relative errors between LO and FE models were small, with a maximum relative error of ~5% observed for the HF geometry at the lowest value of *n*_*A*_. For the final scenario (see rightmost column), we varied *τ*_2_, which impacts the relaxation time of the intracellular calcium transients and, hence, contraction duration. Higher *τ*_2_ values resulted in prolonged ejection and isovolumic relaxation times, and a maximum mismatch of 5% between LO and FE results was observed for the LVV trace of the HF ventricle at the longest considered *τ*_2_ (i.e., for *τ*_2_ = 285 ms).

Overall, the good matches between FE and LO global outputs observed both under training conditions (see Figure 6) and for the simulated scenarios of Figure 7 suggested that, despite accounting for just one active cell unit, the LO models were able to effectively capture the global behavior of 3-D FE models, which instead results from the cooperation of many units operating under heterogeneous strain conditions. For further investigation, we then decided to extend the LO modeling framework to account for heterogeneity by allowing for the global ventricular dynamics to be captured by multiple LO sub-components connected either in parallel or in series and varying for their material properties (see section Multi-element low-order models). More specifically, for units connected in parallel, we imposed variations in *µ*_1_ within the range [*µ*_1,*min*_, *µ*_1,*max*_] = [0.01, 0.179]. Similarly, heterogeneity was achieved for the series configuration cases by modulating 6 LO parameters, {*a, b, c, α, β, µ*_2_}, with weights varying within the range [*w*_*min*_, *w*_*max*_] = [0.6, 1.0]. To analyze the capability of single-element models to approximate multi-element configurations, we then trained LO models with a single cell unit to maximize the match in global ventricular volumes and pressures. Average mismatches, quantified as *C* (see (5)), between single and multi-element models are reported in Table 4 upon variations of the same testing parameters considered for Figure 7. In all cases, the single element models were able to capture well the multi-unit behaviors, with a maximum mismatch of 3.5% observed for the parallel configuration at the lowest *V*_*max*_ considered. Maximum mismatch with the series configuration was observed instead for the largest *τ*_2_ value (*C*=2.59%).

**Table 4.** Error of approximation of multi-element models by the LO model with a single element.

| Parallel |  |  |  |  |  |  |  |
| --- | --- | --- | --- | --- | --- | --- | --- |
| $V_{max}$ | | $n_0$ | | $n_A$ | | $\tau_2$ | |
| $s_v$ factor | error, x100 | value | error, x100 | value | error, x100 | scaling factor | error, x100 |
| 1.00 | 0.53 | 0.50 | 0.62 | 1.00 | 0.11 | 1.10 | 0.54 |
| 1.75 | 0.51 | 0.54 | 0.58 | 2.00 | 0.14 | 1.20 | 0.57 |
| 2.50 | 0.55 | 0.59 | 0.54 | 3.00 | 0.21 | 1.30 | 0.60 |
| 3.25 | 0.53 | 0.63 | 0.53 | 4.00 | 0.51 | 1.40 | 0.63 |
| 4.00 | 0.50 | 0.68 | 0.53 | 5.00 | 0.89 | 1.50 | 0.65 |
| 0.62 | 1.15 | 0.72 | 0.57 | 6.00 | 0.83 | 1.60 | 0.68 |
| 0.45 | 1.82 | 0.77 | 0.66 | 7.00 | 0.75 | 1.70 | 0.70 |
| 0.36 | 2.40 | 0.81 | 1.13 | 8.00 | 0.66 | 1.80 | 0.72 |
| 0.29 | 2.94 | 0.86 | 0.93 | 9.00 | 0.57 | 1.90 | 0.74 |
| 0.25 | 3.43 | 0.90 | 2.08 | 10.00 | 0.53 | 2.00 | 0.75 |
| Series |  |  |  |  |  |  |  |
| $V_{max}$ | | $n_0$ | | $n_A$ | | $\tau_2$ | |
| $s_v$ factor | error, x100 | value | error, x100 | value | error, x100 | scaling factor | error, x100 |
| 1.00 | 1.31 | 0.50 | 1.57 | 1.00 | 2.99 | 1.10 | 1.38 |
| 1.75 | 1.84 | 0.54 | 1.48 | 2.00 | 3.15 | 1.20 | 1.47 |
| 2.50 | 2.26 | 0.59 | 1.40 | 3.00 | 3.47 | 1.30 | 1.57 |
| 3.25 | 2.44 | 0.63 | 1.31 | 4.00 | 2.55 | 1.40 | 1.69 |
| 4.00 | 2.48 | 0.68 | 1.23 | 5.00 | 3.06 | 1.50 | 1.83 |
| 0.62 | 1.19 | 0.72 | 1.15 | 6.00 | 2.58 | 1.60 | 1.98 |
| 0.45 | 1.22 | 0.77 | 1.04 | 7.00 | 2.08 | 1.70 | 2.14 |
| 0.36 | 1.16 | 0.81 | 1.07 | 8.00 | 1.72 | 1.80 | 2.30 |
| 0.29 | 1.16 | 0.86 | 1.46 | 9.00 | 1.47 | 1.90 | 2.45 |
| 0.25 | 1.16 | 0.90 | 1.45 | 10.00 | 1.31 | 2.00 | 2.59 |

As an example, Figure 8 shows pressure (top row) and volume (bottom row) traces from multi-element models (connected in series) overlayed on the corresponding traces from best-fit single-element models. The results shown derived from two different levels of myofilament cooperativity (i.e., *n*_*A*_ = 10 and and *n*_*A*_ = 5, in the left and right panel, respectively) demonstrating almost complete overlap between outputs from the two modeling schemes (i.e., compare dashed and dotted black lines). Heterogeneity within multi-element models was evident when comparing volume traces from the two most different subunits of the model (i.e., those modulated by weights at the range extremes). As weights modulated also passive stiffness, the *w*_*max*_ (*w*_*min*_) subunit was least (most) compliant and therefore contributed the smallest (largest) volume subportion to the resulting total trace. As expected, no differences could be appreciated among pressure traces, as subunits connected in series were designed to sustain the same left ventricular pressure.

**Fig 8.**
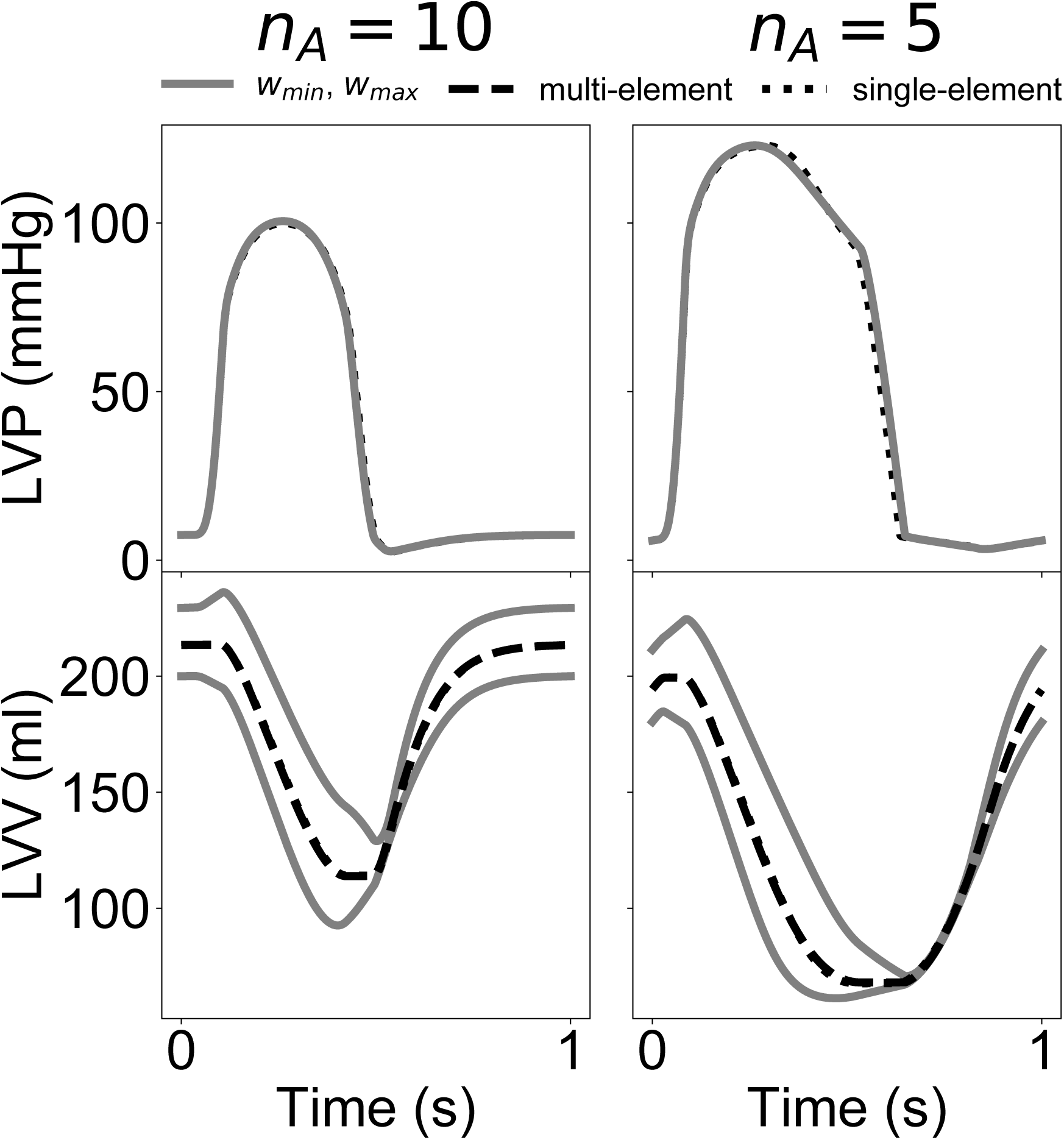
Global output comparison between series multi-element and single-element LO models at two levels of myofilament cooperativity (i.e., *n*_*A*_ = 5 and *n*_*A*_ = 10). Pressure (top), and volume (bottom) traces are shown for the two multi-element subcomponents with extreme parameter weights (i.e., *w*_*min*_ = 0.6 and *w*_*max*_ = 1.0, respectively; see gray lines). Total output traces from the multi-element models (see dashed black lines) are also shown together with analogue output traces from the single-element models (see dotted black lines). Total traces show almost complete overlap.

As the the above results show, after training, even single cell-to-global transformations (2) were able to capture the structural organization and heterogeneity accounted for in FE models and that we imposed in our multi-element LO models. To characterize better the links existing between ventricular anatomy and corresponding trained LO parameters, we then repeated congruency training for a set of virtual ventricles representative of the anatomies collected by the Sunnybrook Cardiac MRI database. Similar to [27], we employed an axisymmetric 6-parameter geometric description and used latin hypercube sampling to extract 150 ventricles from uniform distributions constructed over the entire range of the dataset anatomies. From this initial sampling, 44 simulations were discarded as either not compatible (e.g., due to unrealistically large ventricular thickness as compared to radius), or because they did not contract enough (i.e., LVV did not reach 85% of V_0_ at peak contraction, see section Regression for congruency coefficients for more details). For the remaining 106 simulations, congruency training was achieved with mean square errors always lower than 1%. Figure 9 shows scatter plots and overall trends of the LO parameters after training from the active simulations (*µ*_1_, *µ*_2_ and *γ*) with respect to the variations of the two most influential anatomical parameters (*R*_*b*_ and the ventricular thickness *L*). When considering the trends separately (see panel A), all trained parameters were most affected by *L*. Some of this dependence on L can be expected, as thicker ventricles are able to accomodate more force-generating myocytes, thus explaining the direct proportionality between *L* and *µ*_2_ (see central row). The opposite was true for *µ*_1_, which transforms LVV into myocardial strain, indicating that, in thicker ventricles, changes in volumes tend to correspond to smaller changes in effective myocardial strain (see top row). Effects on *γ* (see bottom row) were more complex with a minimum *γ* = 0.92 observed for *L* = 13 mm and higher values observed both at smaller and larger thicknesses. The role played by *R*_*b*_ emerged more clearly in contour plots showing expected coupled effects when varying also *L* in a 2-D Gaussian Process regression (see panel B). In general, *R*_*b*_ had opposite effects with resepect to *L*, which resulted in the maximum expected parameter values being located in the vicinity of either the top (for *µ*_1_) or bottom (for *µ*_2_ and *γ*) left corners. In addition, the combined variations assumed locally complex behaviors (e.g., see isocontours in the lower half the *µ*_1_ plot), which would difficult to capture without a statistical regression model.

**Fig 9.**
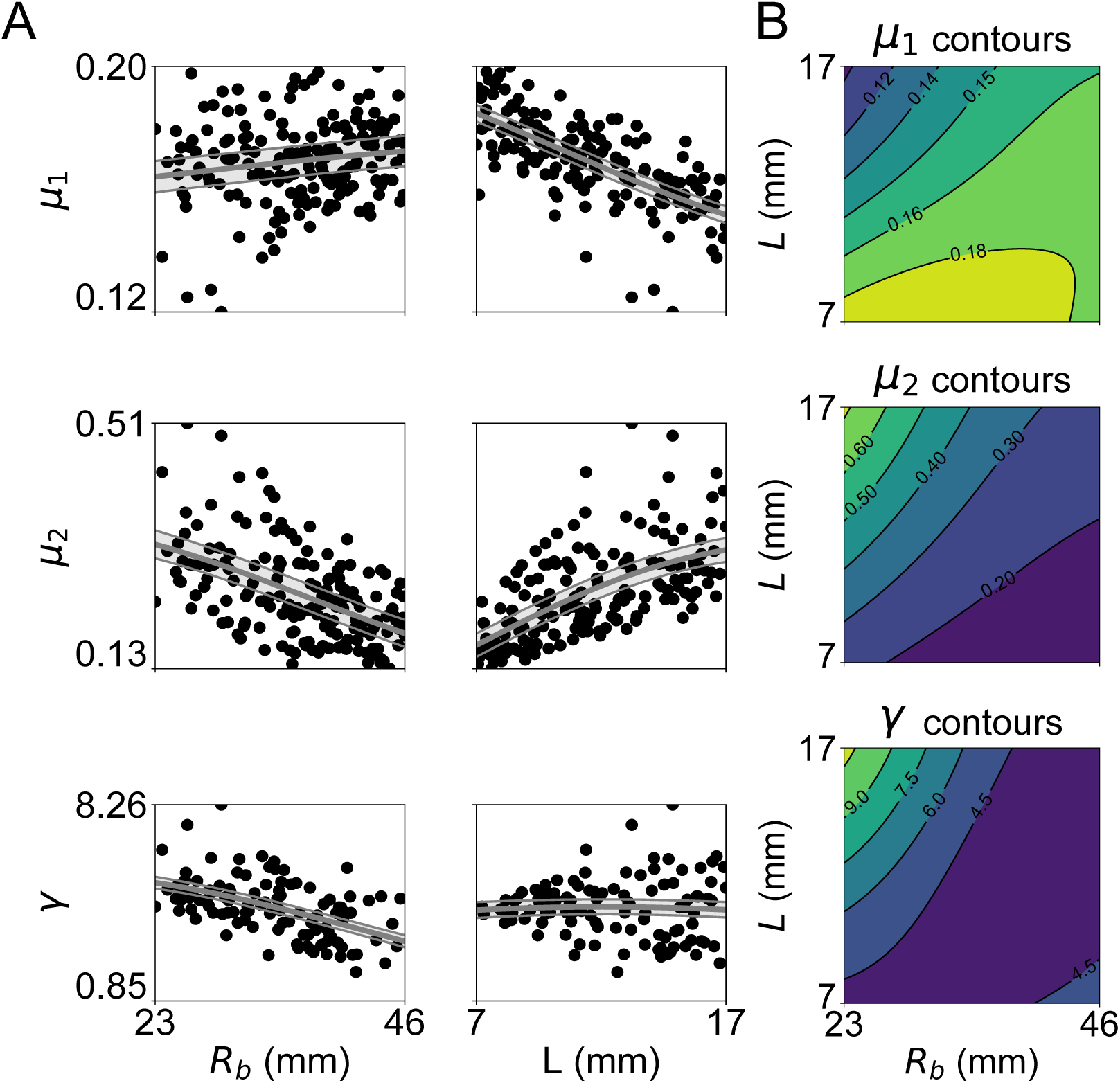
Variations of parameter trained via active simulations (*µ*_1_, top row; *µ*_2_, central row; *γ*, bottom row) with respect to changes in ventricular anatomy. A) Scatter plots showing trends with respect to variations in *R*_*b*_ (left column) and *L* (center column). Grey overlay indicates expected value (solid lines) and standard deviation of a 1-D Gaussian Process regression fitted on the data. B) Contour plots of trained parameter values expected for combined variations in *R*_*b*_ and *L* as predicted by a 2-D Gaussian Process regression model fitted to the data.

Fig 10 shows cross-validation results for the Gaussian Process regression models trained to map the 6 anatomical parameters representing the axisymmetric ventricles to the 8 LO parameters. In total, we constructed 5 regression models on training sets of increasing size (i.e., n = 10, 20, 50, 106) and evaluated 5-fold cross-validation errors for each. In all cases, as expected, larger training sets yielded better accuracy, with errors falling below 3% for all parameters except for *γ*, which could be inferred only with an everage 6% error. The *γ* parameter plays a role only during part of systole and it is therefore not surprising that its estimation was more difficult, as it is less well characterized by our training set. Error was lower for the remaining “trained” parameters affecting active contraction (i.e., 2% and 3% cross-validation errors for *µ*_1_ and *µ*_2_, respectively, see top panel), and always smaller than 2% for the passive material property parameters (see central and bottom panel).

**Fig 10.**
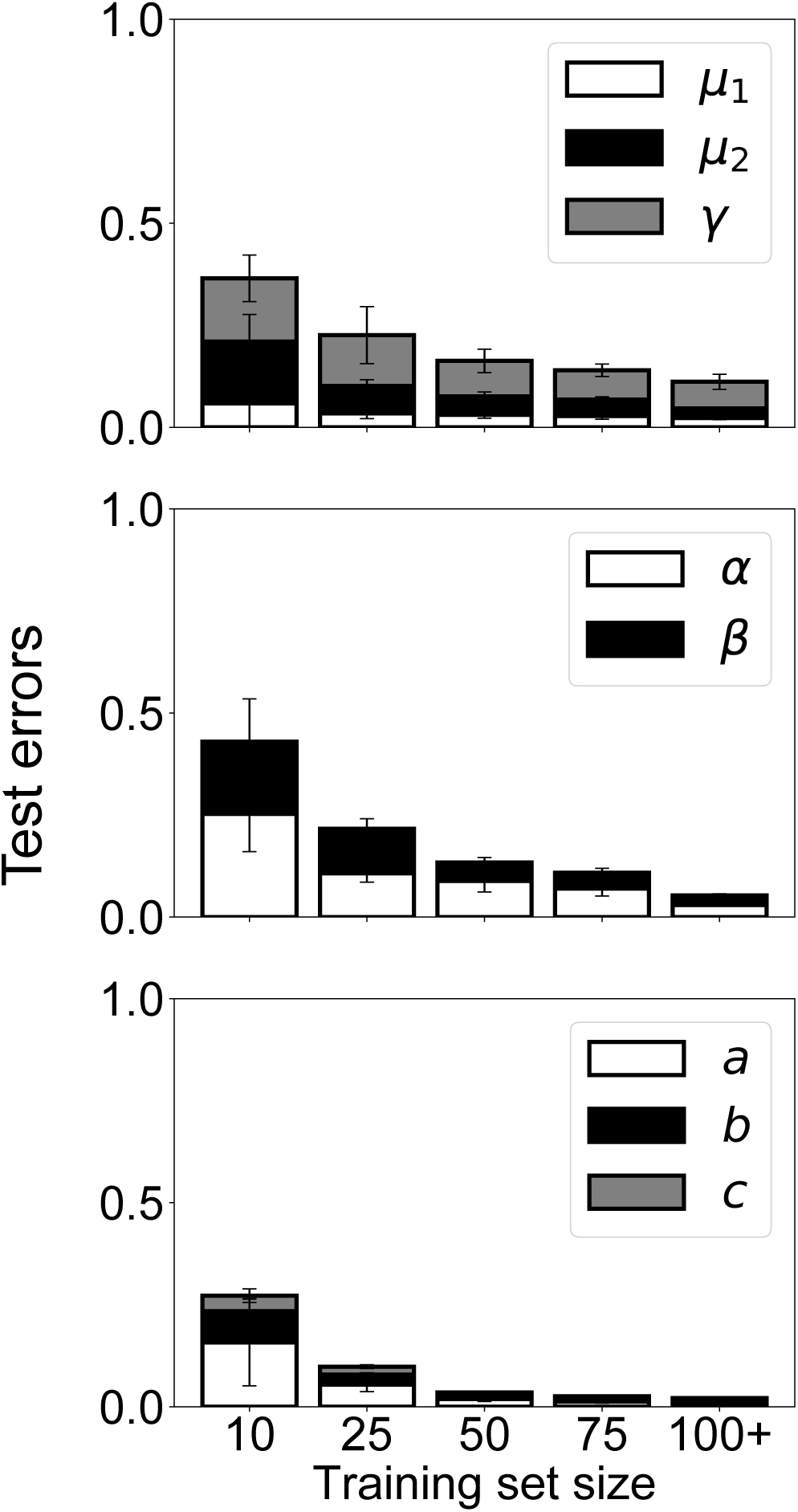
5-fold cross-validation of GP regression models mapping LV geometric features to active and passive LO model parameters.

Figure 11 shows simulated global outputs after the model parameters were optimized to match cardiac phase durations and ejection fraction from MR imaging for the N (left panels) and HF (right panels) cases. In addition, we show volume measurements extracted from MRI that provided the target for the optimization (see solid circles in the two bottom panels). As expected, the MRI analysis revealed a reduced ejection fraction (−19%) and a prolonged ejection time (+21% after normalizing to a 60 bpm heart rate) for the HF case than to the N one (see Table 5). Accordingly, the optimization algorithm achieved optimal matches by assigning smaller *S*_*a*_ (−46%) and larger *τ*_2_ (+31%) values to the HF case compared to the N one. Given our simplified description of the atrium, we did not correctly reproduce the dynamics of diastoling filling. Accordingly, our target variables (i.e., ejection fraction, ejection time, isovolumic relaxation time, and aortic pulse pressure) did not include diastolic filling metrics, as they would not be helpful without proper accounting of atrial and mitral valve function. Average mismatches to the target variables were smaller than 3% in both cases, suggesting that the combined tuning of hemodynamic parameters, calcium handling via *τ*_2_ and contraction efficiency via *S*_*a*_ might be sufficient to constrain the model to global contraction measurements from images.

**Fig 11.**
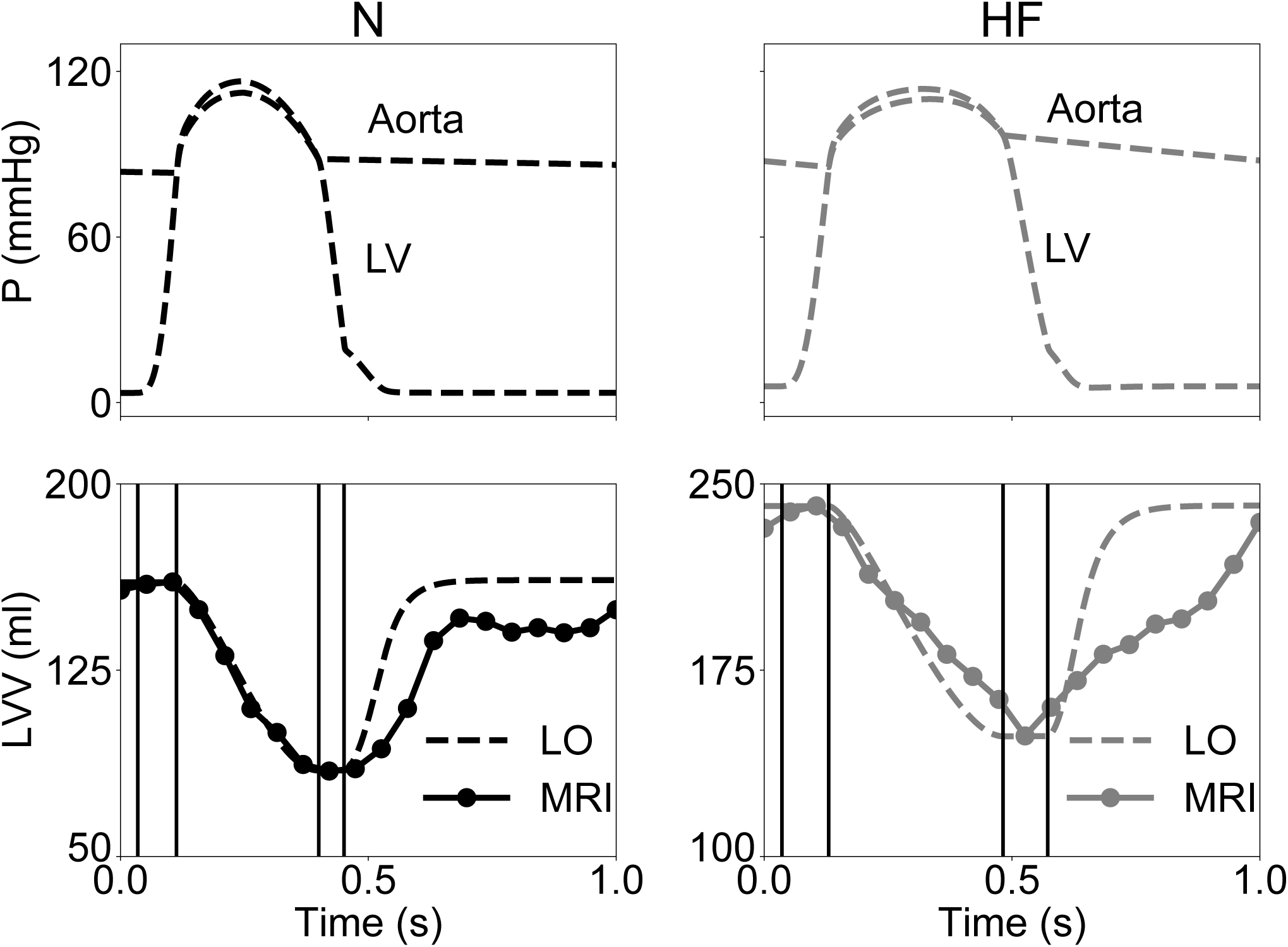
Inverse estimation of model parameters from cardiac phase duration and ejection fraction measured from MRI. Top row, simulated intraventricular and aortic pressure traces after optimization for the N (left) and HF (right) cases. Hemodynamic parameters of the Windkessel submodel were adapted to maintain central aortic pressure approximately within the normal range 80-120 mmHg. Bottom row, corresponding simulated ventricular volume traces (see dashed lines) overlayed to measurements from 20 MRI snapshots (see solid circles) included in the Sunnybrook Cardiac MRI database. The *τ*_2_ and *S*_*a*_ parameters (regulating calcium handling and contraction efficiency, respectively) were optimized to match ejection and isovolumic relaxation times, as well as ejection fraction. Solid vertical lines mark transitions between simulated cardiac phases, denoting good match with corresponding transitions in the MRI traces. As the model does not properly account for atrial and mitral valve functions, behaviors during diastolic filling are are not expected to mach well, and do not.

**Table 5.**
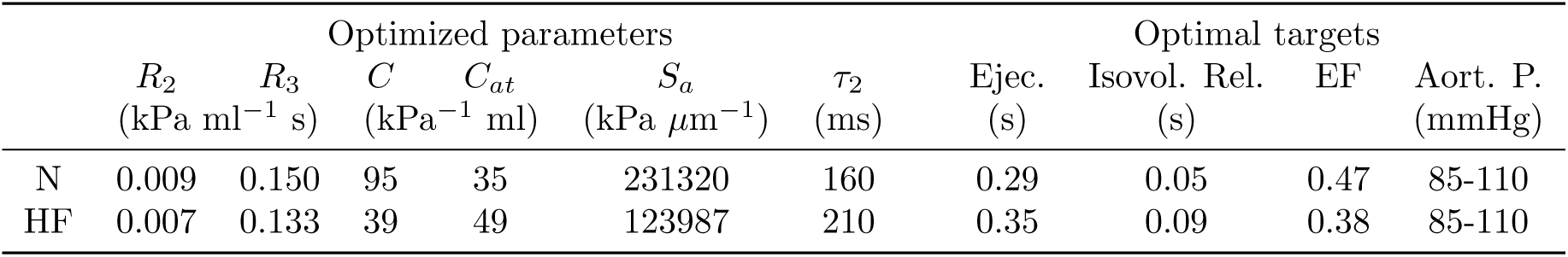
Optimal model parameters estimated to match cardiac phase durations and ejection fraction from MRI

## Discussion

Models of cardiac mechanics promise to provide significant insights into the state of health of patients with heart disease by providing quantitative descriptions of the biophysics driving cardiac contraction at different space and time scales. Image-based mechanistic reconstructions such as state-of-the-art FE models allow for capturing high levels of detail in anatomic complexity and spatial heterogeneity (e.g., to account for the varying preferential directions of myofibers), but are also computationally expensive. Not surprisingly, significant obstacles are added when large parameter searches and inverse problem solutions are required to adapt these models to experimental or clinical observations. To at least partially address such challenges, we have demonstrated a procedure to train efficient LO models of ventricular mechanics and reproduce the global behavior of FE models by scaling the outputs of just a single modeled cell. The computational costs of cardiac models were therefore radically reduced. Our strategy was motivated by our finding that the relationship between local fiber strain and ventricular volume could be well linearized over a large range of stretches (see Fig 5), and surprisingly also when accounting for material and geometric nonlinearities [33, 34], and spatially-varying distribution of fiber orientation and heterogeneous activation times [35]. By lumping the nonlinearities introduced by the 3-D architecture of the myocardial tissue at the passive level (e.g., during filling) into an additive pressure term (see (1)), we were able to reproduce the global behaviors observed at the organ level by just linearly scaling the active contributions of the cellular model (see (3)). In effect, our approach exploited the links between length dependence active force and Frank-Starling mechanism [38] to generate a concise model of global heart function, which can be computed with minimal cost. Coupling between local cell behavior and global outputs was achieved by transformation modules incorporating the linear relationships (3), which were trained to ensure congruency with FE results upon varying hemodynamic conditions. Given the empirical nature of the transformation relationships, the “trained” parameters {*µ*_1_, *µ*_2_, *γ, α, β, a, b, c*} did provide only an indirect physiological meaning. Although we did not conduct a systematic investigation to minimize the number of high-resolution simulations necessary for training, our results suggested that even just 3 active and one passive high-resolution simulations were sufficient to achieve good fits of the LO model to the FE model (see Fig 6).

We submit that the presented model order reduction strategy presents significant advantages over standard machine learning approaches where training is conducted directly on high-resolution simulations (e.g., [27, 31]). While efficient at the inference stage, statistical models tend to require extensive training to achieve sufficient accuracy. By maintaining a biophyisical component (i.e., the cell unit), our LO models can be more easily trained, especially given that probing simulations can be chosen to explore large variations in operating sarcomere lengths (see Fig 5 and Table 2). After training, LO models then achieve remarkable computational performance, with the possibility of simulating thousands of heartbeats every second on commodity hardware. This level of computational efficiency is comparable to alternative order reduction methods [13–16], but, unlike these, our approach does not require restrictive assumptions on ventricular anatomy. The popular CircAdapt model [13], for example, achieves computational efficiency by assuming sphericity of ventricles, and then by analytically exploiting symmetry. The congruency training proposed here can handle any anatomy that the FE model can handle. The LO model also outperform the speed-ups typically reported for more sophisticated numerical approaches, such as reduced-basis functions and proper orthogonal decomposition (e.g., [17, 18, 21, 22]), although, in the current form, our reduction technique does not maintain details on spatially-varying fields.

To investigate further the properties of the LO reduction approach, we also considered extensions of the single-element LO architectures, and built configurations where multiple LO subcomponents are connected either in series or in parallel (see Fig 4). Even when introducing heterogeneity among material properties of different subcomponents, the global outputs could still be recapitulated by a trained single-cell analogue. This finding was also backed analytically under the assumptions of a simplified model of active contraction and of linear passive material properties (see section “Approximation of multi-element model with single-element LO model” in the Supplemental Material). Our empirical and analytical analyses demonstrate that the global aspects of the biomechanics simulated by FE models could indeed be captured by these simplified descriptions.

Given their computational efficiency and demonstrated versatility, the presented LO models are thus ideal for solving difficult inverse parameter estimations problems. To showcase this strategy, we adapted model parameters to capture contraction features measurable from cardiac imaging. More specifically, we relied on our previous analysis of the Sunnybrook Cardiac MRI database [27] to select 2 representative patients from the normal and heart failure groups, and found model parameterizations that would match ejection fraction and cardiac phase durations from MRI, while also maintaining central aortic pressure within a reasonable 85-110 mmHg range. As expected for this type of complex inverse estimation problem, parameter optimization required running the LO models for several thousands of parameter combinations and for tens of beats at each evaluation to reach steady-state. Tackling such an inverse problem would be impractical with a standard FE technique, even when considering efficient surrogate-based optimization algorithms [28, 29, 37], as we did in this study. By exploiting multi-core execution, we were instead able to optimize LO model parameters in just a few minutes on commodity hardware, and achieved good matches (see Fig 11). Interestingly, the optimization algorithm did not reveal large differences in hemodynamic lumped parameters, but it attributed instead the discrepancies in ejection fraction and cardiac phase duration to the *S*_*a*_ and *τ*_2_ parameters, which modulate cardiac contraction efficiency and decay time of calcium transients, respectively. More specifically, our analysis suggested that the HF MRI measurements could be explained by smaller *S*_*a*_ values and longer *τ*_2_ times, which is in agreement with experimental findings on heart failure myocyte phenotypes, which are less effective in contracting partially due to dysfunctional calcium handling [39, 40].

Given the encouraging preliminary results in extracting mechanistic insights from clinical assessments of cardiac function, we submit that, after validation, our approach could prove helpful to assist statistical analyses of large databases of clinical measurements and provide interpretations for variable associations. By providing an efficient model-order reduction strategy that does not compromise anatomical details, our work might also facilitate future studies involving high resolution simulations, where FE and LO models could be employed in combination for effective inverse estimation of parameter values.

## Acknowledgments

Part of this work was carried out within the framework of the IIF UrB RAS government assignment and was partially supported by the UrFU Competitiveness Enhancement Program (agreement 02.A03.21.0006) as well as the RSF grant (No. 19-14-00134). The URAN supercomputer at IMM UrB RAS was used for part of the model calculations.

## Conflict of interest

Paolo Di Achille, Jaimit Parikh, James Kozloski and Viatcheslav Gurev are employees of IBM.

## Notes

#### Summary of Updates

The supplemental material was added.

