## Supplemental material for "Model order reduction for left ventricular mechanics via congruency training"

### Myofilament contraction model

The LO simulations make it possible to design novel cardiac muscle contraction models by following iterative approaches where the consequences of model assumptions can be evaluated across scales from isolated preparation assays to whole organ effects. Leveraging such capability, we constructed a new phenomenological model of muscle contraction. The new myofilament model is a modified version of [1] with the addition of XB-XB cooperative effects and with a simple mean-field strain formulation similar to [2]. The model equations are designed to reproduce several experimental features observed in both isolated muscle and physiological measurements at the ventricle level, as described below.

#### XB kinetics and ktr-Ca dependence

Similar to previous models describing cooperative behavior of crossbridges (XBs) within regulatory units (RUs) (e.g., [3]), we simulated XB dynamics from the perspective of XB-groups. The core model equation describes the formation and collapse of XB "populations", groups of XBs featuring XB-XB cooperative effects within a group. The XB-group could be interpreted as the crossbridges in a single RU, as crossbridges probably interact only within a RU [4]. Alternatively, groups could be interpreted as all RUs of separate actin filaments, if we were to assume that all crossbridges in the actin-activation model interact with each other. Finally, populations could be purely functional units.

In a typical XB-group development, force is first generated when a "primary" crossbridge is formed as a myosin head binds actin at a relatively slow rate, initiating a new group. "Secondary" crossbridges are then formed at a much faster rate, which results in collapsing of groups at relatively slow rates. The simplest equation for the fraction of groups $G_{XB}$ engaged in force generation that guarantees a mono-exponential shape for the rate of force redevelopment ($Ktr$) curve assumes the form

$$\frac{d\mathrm{G}_{XB}}{dt} = f_G(A)(1 - \mathrm{G}_{XB}) - g_G(A, s)\ \mathrm{G}_{XB}, \tag{1}$$

where $f_G$ and $g_G$ rates are both functions of the fractions of troponin complexes with bound Ca ($A$), and $g_G$ is also a function of the mean XB strain ($s$). We assumed that all XB-groups have the same size, and that XB-group size depends on the average rates of myosin binding and detachment within the XB-group. Then the fraction of bound XBs ($\mathrm{XB}_G$) within a group can be described by

$$\frac{d\mathrm{XB}_G}{dt} = f_{XB}(1 - \mathrm{XB}_G) - g_{XB}(s)\ \mathrm{XB}_G, \tag{2}$$

where the rate $f_{XB}$ is a constant, and $g_{XB}$ is a function of the XB strain. We can assume that the group formation rate $f_G$ is mostly determined by the slow binding rate for formation of primary XBs, while formation of secondary XBs within the same group follows a much faster process. The total fraction of crossbridges in the force-generating state is equal to $G_{XB} \times XB_G$.

To ensure calcium dependence of $Ktr$, the rates $f_G$ and $g_G$ were formulated as functions of $A$, giving a nearly Hill-shaped curve for the force-calcium relationship,

$$\begin{aligned} f_G\left(A\right) &= \bar{f}_G \left( \frac{A^{n_A}}{A^{n_A} + A_{50}^{n_A}} \right)^{\zeta} \\ g_G\left(A, s\right) &= max \left\{ g_G(s) \left( \frac{A^{n_A}}{A^{n_A} + A_{50}^{n_A}} \right)^{\zeta - 1}, g_{Gmax} \right\}, \end{aligned} \tag{3}$$

where $0 \leq \zeta \leq 1$ and $\bar{f}_G$ are model parameters, $g_G$ is a function of XB strain, and $A_{50}$ and $n_A$ are parameters that define cooperativity in the interaction between troponin, tropomyosin state, and crossbridge dynamics in the XB-groups.

#### XB strain

The equation for the mean XB strain $s$ was derived using a 2-state scheme for XBs, derived as in [2]

$$\frac{ds}{dt} = \frac{1}{2}\frac{dSL}{dt} - f_{XB}\frac{1 - \mathrm{XB}_G}{\mathrm{XB}_G}s, \tag{4}$$

where $SL$ is the sarcomere length. The rates $g_{XB}$ and $g_G$ in (2) and (3) are functions of $s$. We introduced a factor $s_{mod}\left(s\right) = e^{-\alpha(s/x_0)^2}$ to modify the rates

$$\begin{aligned} g_{XB}\left(s\right) &= \bar{g}_{XB} \times s_{mod}\left(s\right) \\ g_G\left(s\right) &= \bar{g}_G \times s_{mod}^{\eta}\left(s\right), \end{aligned} \tag{5}$$

where $x_0$ is the distortion due to XB power-stroke, $\bar{g}_G$, $\bar{g}_{XB}$, $\alpha$, and $\eta$ are parameters of the model.

#### Calcium binding and length dependent activation

We maintained the same functional form of length dependence for the thick filament as in [1] by using a piecewise polynomial function. Following the max-min representation of piecewise linear functions [5], we employed the form

$$\mathrm{LDF}_{\mathrm{thick}}\left(\lambda\right) = 0 \vee \left[\mathrm{LDF}_{\mathrm{thickmax}} \wedge \left(\lambda_{\mathrm{ms0}} \times \left(\lambda - \lambda_{\mathrm{mn0}}\right)\right) \wedge \left(\lambda_{\mathrm{ms1}} \times \left(\lambda - \lambda_{\mathrm{mn1}}\right) + \mathrm{LDF}_{\mathrm{thickmax}}\right)\right], \tag{6}$$

where $\wedge$ and $\vee$ are **min** and **max** binary operators, respectively, $\lambda$ is the stretch ratio of a sarcomere (we assume a sarcomere length of 1.9 $\mu m$ at $\lambda = 1$), $\lambda_{\mathrm{ms0}} > \lambda_{\mathrm{ms1}}$ are the slopes of the piecewise linear function, and $\lambda_{\mathrm{mn0}} < \lambda_{\mathrm{mn1}}$ are the nodes defining the function discontinuities. The length dependend function (LDF) has a minimum 0 and a maximum

$\mathrm{LDF}_{\mathrm{thickmax}}$. The active tension developed by the muscle ($T_a$) is then calculated as

$$T_a = S_a \times \mathrm{LDF}_{\mathrm{thick}}(\lambda) \times G_{XB} \times XB_G \times (s + x_0), \tag{7}$$

where $T_a$ is the active tension; $S_a$ is the scaling factor for the tension; $x_0$ is the step size of the myosin power stroke.

Cardiac troponin complexes modulate myofilament contraction in a Ca-dependent manner by regulating the availability of myosin binding sites on actin. As in [1], regulation of high and low affinity troponin was captured by two separate equations

$$\frac{d}{dt}\mathrm{A}_H = k_{\mathrm{onT}}[\mathrm{Ca}](1 - \mathrm{A_H}) - k_{\mathrm{offHT}}\mathrm{A}_H, \tag{8}$$

$$\frac{d}{dt}\mathrm{A}_L = k_{\mathrm{onT}}[\mathrm{Ca}](1 - \mathrm{A}_L) - k_{\mathrm{offLT}}\mathrm{A}_L, \tag{9}$$

$$\mathrm{A} = \mathrm{A}_H \times \mathrm{LDF}_{\mathrm{thin}}(\lambda) + \mathrm{A}_L \times (1 - \mathrm{LDF}_{\mathrm{thin}}(\lambda)), \tag{10}$$

where $\mathrm{A}_H$ and $\mathrm{A}_L$ refer to the fraction of high and low calcium affinity troponin sites, respectively, that have Ca bound to their regulatory binding sites; $k_{\mathrm{onT}}$ is the rate constant for binding; [Ca] is the concentration of Ca; $k_{\mathrm{offHT}}$ and $k_{\mathrm{offLT}}$ are the rate constants for unbinding from the high and low affinity sites, respectively. As for the thick filament, a function for length dependence is defined as

$$\mathrm{LDF}_{\mathrm{thin}}(\lambda) = 0 \vee \left[\mathrm{LDF}_{\mathrm{thinmax}} \wedge (\lambda_{\mathrm{as0}} \times (\lambda - \lambda_{\mathrm{an0}})) \wedge (\lambda_{\mathrm{as1}} \times (\lambda - \lambda_{\mathrm{an1}}) + \mathrm{LDF}_{\mathrm{thickmax}})\right]. \tag{11}$$

The equation for intracellular calcium was taken from [1]

$$\beta = \left(\frac{\tau_1}{\tau_2}\right)^{-1/\left(\frac{\tau_1}{\tau_2}-1\right)} - \left(\frac{\tau_1}{\tau_2}\right)^{-1/\left(\frac{\tau_2}{\tau_1}-1\right)}, \tag{12}$$

$$[Ca](t) = \begin{cases} Ca_{diast},\ t \le t_{start} \\ \left(\frac{Ca_{amp} - Ca_{diast}}{\beta}\right) \times \left(e^{-\frac{t - t_{start}}{\tau_1}} - e^{-\frac{t - t_{start}}{\tau_2}}\right) + Ca_{diast},\ t > t_{start} \end{cases}. \tag{13}$$

### Summary of model equations

$$
\begin{aligned}
&\beta = \left(\frac{\tau_1}{\tau_2}\right)^{-1/\left(\frac{\tau_1}{\tau_2}-1\right)} - \left(\frac{\tau_1}{\tau_2}\right)^{-1/\left(\frac{\tau_2}{\tau_1}-1\right)} \\
&[Ca]\,(t) = \begin{cases} Ca_d,\ t \le t_{start} \\ \left(\frac{Ca_{amp}-Ca_{diast}}{\beta}\right) \times \left(e^{-\frac{t-t_{start}}{\tau_1}} - e^{-\frac{t-t_{start}}{\tau_2}}\right) + Ca_{diast},\ t > t_{start} \end{cases} \\
&SL = 1.9\mu m \times \lambda \\
&s_{mod} = e^{-\alpha (s/x_0)^2} \\
&f_G\,(A) = \bar{f}_G \left(\frac{A^{n_A}}{A^{n_A} + A_{50}^{n_A}}\right)^{\zeta} \\
&g_G\,(s) = \bar{g}_G \times s_{mod}^{\eta} \\
&g_{XB}\,(s) = \bar{g}_{XB} \times s_{mod} \\
&g_G\,(A, s) = max\left\{ g_G(s) \left(\frac{A^{n_A}}{A^{n_A} + A_{50}^{n_A}}\right)^{\zeta - 1}, g_{Gmax} \right\} \\
&\frac{d\mathrm{G}_{XB}}{dt} = f_G(A)(1 - \mathrm{G}_{XB}) - g_G(A,s)\ \mathrm{G}_{XB} \\
&\frac{d\mathrm{XB}_G}{dt} = f_{XB}(1 - \mathrm{XB}_G) - g_{XB}(s)\ \mathrm{XB}_G \\
&\frac{ds}{dt} = \frac{1}{2}\frac{dSL}{dt} - f_{XB}\frac{1 - \mathrm{XB}_G}{\mathrm{XB}_G} s \\
&\frac{d}{dt}\mathrm{A}_H = k_{\mathrm{onT}}[\mathrm{Ca}](1 - \mathrm{A_H}) - k_{\mathrm{offHT}}\mathrm{A}_H \\
&\frac{d}{dt}\mathrm{A}_L = k_{\mathrm{onT}}[\mathrm{Ca}](1 - \mathrm{A}_L) - k_{\mathrm{offLT}}\mathrm{A}_L \\
&\mathrm{A} = \mathrm{A}_H \times \mathrm{LDF_{thin}}\,(\lambda) + \mathrm{A}_L \times (1 - \mathrm{LDF_{thin}}\,(\lambda)) \\
&\mathrm{LDF_{thick}}\,(\lambda) = 0 \vee [\mathrm{LDF_{thickmax}} \wedge (\lambda_{\mathrm{ms0}} \times (\lambda - \lambda_{\mathrm{mn0}})) \wedge (\lambda_{\mathrm{ms1}} \times (\lambda - \lambda_{\mathrm{mn1}}) + \mathrm{LDF_{thickmax}})] \\
&\mathrm{LDF_{thin}}\,(\lambda) = 0 \vee [\mathrm{LDF_{thinmax}} \wedge (\lambda_{\mathrm{as0}} \times (\lambda - \lambda_{\mathrm{an0}})) \wedge (\lambda_{\mathrm{as1}} \times (\lambda - \lambda_{\mathrm{an1}}) + \mathrm{LDF_{thinmax}})] \\
&T_a = S_a \times \mathrm{LDF_{thick}}\,(\lambda) \times G_{XB} \times XB_G \times (s + x_0)
\end{aligned}
$$

### Myofilament model simulations

The myofilament model parameters are given in Table 1. The model reproduces several *in vitro* experimental tests (Figure 1) that are typically carried out to characterize myofilament properties. Model simulation of some of these tests is described briefly in the following sections.

#### Isometric twitches

Figure 1A shows simulated force traces obtained under isometric conditions (i.e., fixed sarcomere lengths and transient increases in Ca). Commonly studied characteristics of isometric contraction twitches are the time-to-peak of tension (TPT) and the time from peak of tension to full relaxation ($TR$).

Isometric twitches have roughly the same activation and relaxation rates, so that the twitch shape resembles an

| Parameter | Value (Optimized) | Units |
|---|---|---|
| $\lambda_{\text{an0}}$,$\lambda_{mn0}$ | 0.631579 | unitless |
| $\lambda_{\text{an1}}$,$\lambda_{mn1}$ | 1.21053 | unitless |
| $\lambda_{\text{as0}}$ | 1.58333 | unitless |
| $\lambda_{\text{as1}}$ | 0.791667 | unitless |
| $\text{LDF}_{\text{thickmax}}$ | 1 | unitless |
| $\lambda_{\text{ms0}}$ | 2.45161 | unitless |
| $\lambda_{\text{ms1}}$ | 1.22581 | unitless |
| $\text{LDF}_{\text{thinmax}}$ | 0.645833 | unitless |
| $k_{\text{onT}}$ | 12.5 | $\mu M^{-1}s^{-1}$ |
| $k_{\text{offHT}}$ | 6.25 | $s^{-1}$ |
| $k_{\text{offLT}}$ | 31 | $s^{-1}$ |
| $A_{50}$ | 0.32 | unitless |
| $n_A$ | 10 | unitless |
| $f_{XB}$ | 26.25 | $s^{-1}$ |
| $\bar{g}_{XB}$ | 105 | $s^{-1}$ |
| $\bar{f}_G$ | 25 | $s^{-1}$ |
| $\bar{g}_G$ | 0.07 | $s^{-1}$ |
| $\eta$ | 3 | unitless |
| $\zeta$ | 0.5 | unitless |
| $g_{Gmax}$ | 5e+6 | $s^{-1}$ |
| $Ca_{amp}$ | 0.47 | $\mu M$ |
| $\tau_1$ | 23.07 | ms |
| $\tau_2$ | 190.20 | ms |
| $Ca_{diast}$ | 0.008 | $\mu M$ |
| $\alpha$ | 0.05 (positive strain)<br>1 (negative strain) | |

**Table 1.** Myofilament model parameters

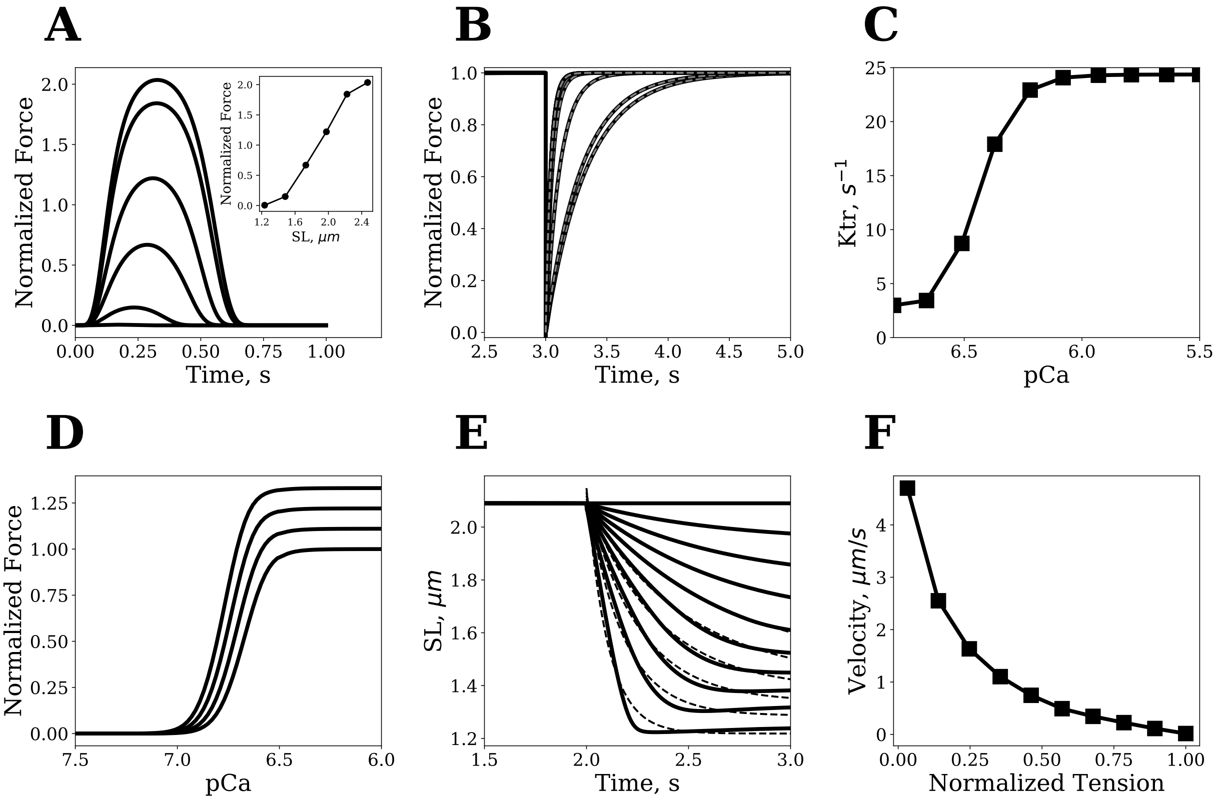


**Fig 1.** *Simulation results from the myofilament model.* ***A****) Force traces during isosarcometric contraction for human cardiomyocytes at 37° C (solid line) for sarcomere lengths (SL) equal to 1.23, 1.48, 1.72, 1.97, 2.22, and 2.47 μm normalized by the peak of force at slack length. The inset shows relationship between force peaks and sarcomere length.* ***B****) Force redevelopment after release-stretch at different Ca level (solid lines) fitted by mono-exponential curve (dashed-line).* ***C****) Rate of force redevelopment at different calcium levels.* ***D****) Force-calcium relationships at different sarcomere lengths.* ***E****) Instantaneous reduction of load on muscle at full calcium activation to register force-velocity curve in* ***F****. Curves after release are fitted with $a + b \times e^{-c(t-t_0)}$, where $t_0$ time of muscle release. Velocity is calculated as a derivative of exponential curve at $t_0$ $(b \times c)$.* ***F****) Simulated force-velocity relationship curve in the model.*

isosceles triangle. In the optimized model, the TPT varied from 303 ms to 326 ms and the time to 95 % relaxation $TR_{95}$ varied from 291 ms to 335 ms for increasing sarcomere lengths (1.9 $\mu m$ - 2.29 $\mu m$). The accelerated relaxation at shorter sarcomere lengths is in agreement with previous experimental observations [6, 7].

#### Rate of force redevelopment $Ktr$

Another common *in vitro* assay for muscle characterization measures involves measuring $Ktr$, the rate of force redevelopment after a sudden length change. To simulate the $Ktr$ protocol, first a steady state force response is achieved at fixed calcium concentration and sarcomere length (2.09 $\mu m$) in the model (as shown in Figure 1B). Next, the sarcomere is released to a lower length, re-stretched and the rate of force redevelopment is examined. The force redevelopment phase is well fitted by a single exponential,

$$P = P_0(1 - e^{-Ktr \cdot t}) \tag{14}$$

where $P_0$ is the initial muscle tension before length-step, and $t$ is time.

Model simulations produce $Ktr$ values on the order of 25 $s^{-1}$, consistent with recordings for human cardiac preparations [8] (see Figure 1C). Moreover, unlike several existing models that fail to reproduce the steep increases of $Ktr$ between low and high Ca concentrations observed experimentally [9], our simulations captured this critical behavior. Simulated $Ktr$ values varied from 25 $s^{-1}$ to 5 $s^{-1}$ for pCa within the 5.6 - 6.4 range. In addition, the model captured the shift of the $Ktr$-pCa relationship to the right of the force-pCa relationship [10].

#### Steady state force-calcium relationship

Figure 1D shows simulated force-calcium (F-Ca) traces obtained at fixed sarcomere lengths (1.9 -2.28 $\mu m$) under constant [Ca]. Model simulations show the typical cooperative relationship between force and calcium concentrations (i.e., significant increase in maximum force for small changes in calcium levels). The simulations also captured the experimentally observed [11] increases in maximal steady force and calcium sensitivity at higher sarcomere lengths. The F-Ca responses are well fitted by a Hill function. The steepness of the F-Ca curve as quantified by the Hill coefficient shows negligible length-dependent effects.

#### Force-velocity relationship

Figure 1E shows observed changes in sarcomere length against different fixed loads at constant calcium concentrations (0.5 $\mu M$). Initially, the muscle is held at a fixed length (SL = 2.08 $\mu m$) and then the load on the muscle is abruptly reduced to lower levels. The velocity of myofilament shortening is obtained through fitting exponential curves to changes in sarcomere length. The model captures the typical inverse relationship between shortening velocity and load (Figure 1F) that is observed experimentally.

#### Physiologic contractions

Physiologic loading conditions are designed to mimic the loads encountered by the muscle during an entire heart cycle. Physiologic contractions have several phases: isometric contraction (which corresponds to isovolumic contraction of the left ventricle), isotonic contraction (ejection phase), isometric relaxation (isovolumic relaxation of the ventricle), and lengthening back to original length with constant velocity (filling phase). In simulations, the filling phase is generated by prescribing a fixed lengthening trajectory. Figure 2A and 2B show simulated force and sarcomere length traces under physiologic loading conditions at different afterloads. The physiologic mode can be used to construct tension-length loops that correspond to pressure-volume loops in whole ventricles (Figure 2C). In agreement with experiments [12, 13], model simulations revealed a linear relationship between tension-length area (a.k.a. stress-strain area) and ATP consumption during a heartbeat (Figure 2D).

### Parameter variation in validation of LO models

As a further validation, we tested optimized LO models upon different sets of conditions that were not explored during congruency training (e.g., see Figure 7 in the result section). In addition, we also trained the single-cell LO model to reproduce outputs from multi-element LO configurations where a certain degree of heterogeneity was introduced by smoothly altering material properties. The first parameter variation that we introduced was in $n_0$, which regulates the

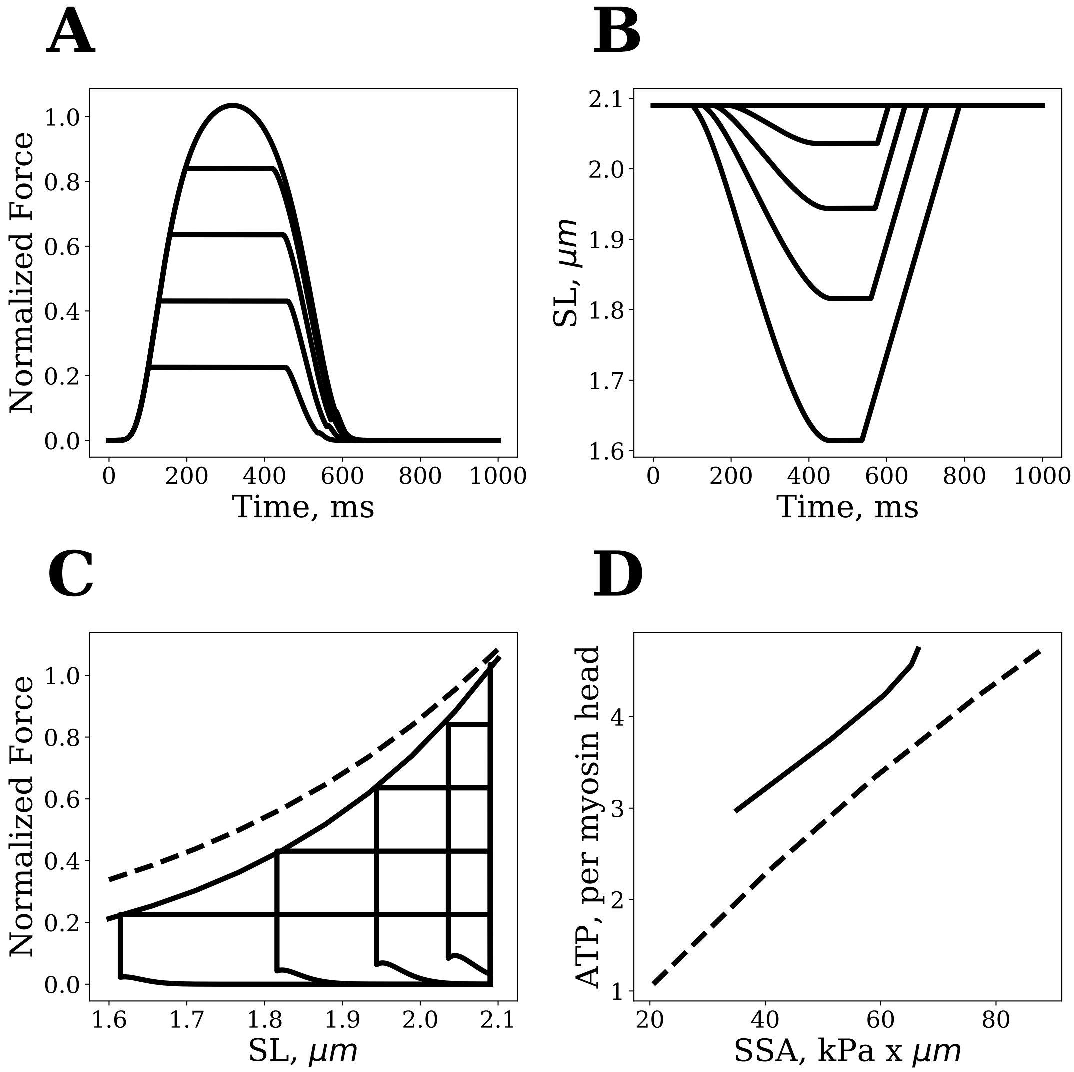


**Fig 2.** *Physiologic contractions (isometric contraction followed by isotonic contraction followed by relengthening)* ***A****. Force traces during physiologic contraction* ***B****. Changes in sarcomere length that correspond to A.* ***C****. Tension-SL loops obtained from A and B during physiologic contraction (analogs to pressure-volume loops for the left ventricle).* ***D****. Stress-strain area vs. ATP consumption for physiologic contraction (black line) and isometric contraction (gray line).*

slope of length-dependence functions according to the following equations (see also Figure 3), 102

$$\begin{aligned}
\lambda_{\mathbf{an0}} &= n_0, \\
\lambda_{\mathbf{mn0}} &= n_0, \\
\Delta &= (\lambda_{an1} - \lambda_{an0}) \,/\, (\lambda_{an1} - n_0)\,, \\
\lambda_{\mathbf{as0}} &= \Delta \times \lambda_{as0}, \\
\lambda_{\mathbf{as1}} &= \Delta \times \lambda_{as1}, \\
\lambda_{\mathbf{ms0}} &= \Delta \times \lambda_{ms0}, \\
\lambda_{\mathbf{ms1}} &= \Delta \times \lambda_{ms1}.
\end{aligned} \tag{15}$$

where the bold font indicates updated values for the considered variables, while normal font weight denotes original values as reported in Table 1. For additional simulations, we also varied the $\tau_2$ and $n_A$ parameters, as well as simultaneously scaled the $f_{XB}$ and $\bar{g}_{XB}$ parameters to vary $V_{max}$ while maintaining similar maximum isometric tension, namely

$$\begin{aligned} \mathbf{f_{XB}} &= s_v \times f_{XB}, \\ \bar{\mathbf{g}}_{\mathbf{XB}} &= s_v \times \bar{g}_{XB}. \end{aligned} \tag{16}$$

Examples of parameter variation effects are shown in Figure 3.

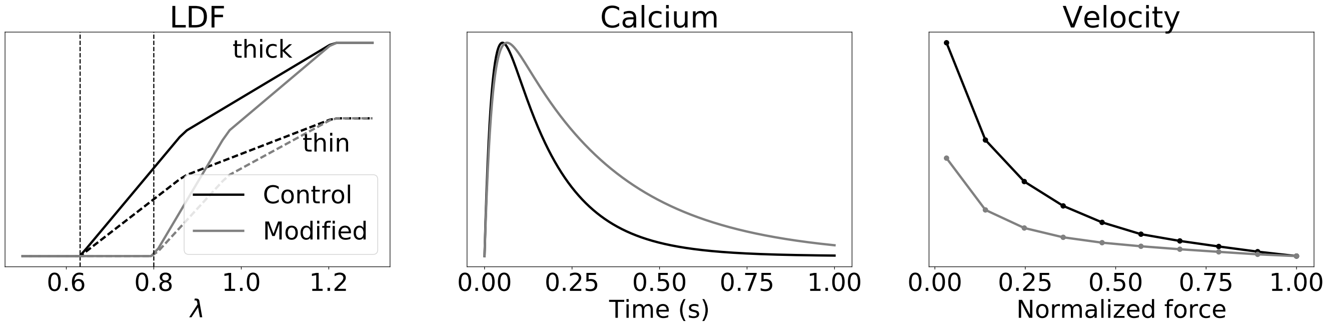


**Fig 3.** *Effects of parameter variation on length dependent functions, calcium transient, and contraction velocity. Control traces and traces after changes of the parameters are shown. LDF is shown for $n_0 = 0.63$ (control) and $n_0 = 0.8$. Calcium transient is shown for $\tau_2 = 153$ (control) and $\tau_2 = 306$. Force-velocity curves are shown for $s_v = 1$ (control) and $s_v = 0.5$.*

### Passive material properties

Passive behavior resulting from the complex architectural organization of myocardial fibers was modeled according to the Fung-type strain energy function used in [14] to model experiments on canine tissue, namely

$$W_U = \frac{C}{2}\left(\exp(Q) - 1\right), \qquad \begin{aligned} Q = {} & b_{\mathrm{ff}} E_{\mathrm{ff}}^2 + b_{\mathrm{ss}} E_{\mathrm{ss}}^2 + b_{\mathrm{nn}} E_{\mathrm{nn}}^2 + b_{\mathrm{fs}}\left(E_{\mathrm{fs}}^2 + E_{\mathrm{sf}}^2\right) \\ & + b_{\mathrm{fn}}\left(E_{\mathrm{fn}}^2 + E_{\mathrm{nf}}^2\right) + b_{\mathrm{ns}}\left(E_{\mathrm{ns}}^2 + E_{\mathrm{sn}}^2\right), \end{aligned} \tag{17}$$

where $E_{\mathrm{ij}}$ $(\mathrm{i}, \mathrm{j} \in \{\mathrm{f}, \mathrm{s}, \mathrm{n}\})$ are components of the Green-Lagrange strain tensor relative to the spatially-varying directions of the myocardial fibers (f), of the myocardial sheets (s) and of the remaining mutually normal direction (n); and $b_{\mathrm{ij}}$ $(\mathrm{i}, \mathrm{j} \in \{\mathrm{f}, \mathrm{s}, \mathrm{n}\})$ are the material property coefficients (values reported in Table 2) expressed in the local reference systems defined by the myocardial fibers and sheets.

**Table 2.** Values for the material property coefficients of (17)

| | Reference | C (kPa) | $b_{\mathrm{ff}}$ | $b_{\mathrm{ss}}$ | $b_{\mathrm{fs}}$ | $b_{\mathrm{n[n,f,s]}}$ |
|---|---|---|---|---|---|---|
| $W_U$ | [14] | 0.88 | 8.00 | 6.00 | 12.00 | 3.00 |

### Approximation of multi-element model with single-element LO model

In this section, we analyze the error made when approximating multi-element (ME) models connected in parallel or series by a single-element LO model. To make the anlysis more tractable, we restricted our focus to cases where active contraction is driven by an elastance-resistance (ER) model. The analysis demonstrates how the LO model resembles in structure and properties the ER model. The LV volume functional, $V$, at a given pressure, $P(t,V)$, is described by a relationship similar to the type 4 model described in [15], namely

$$P(t,V) = k_p(V(t) - V_{p0}) + P_{iso}(t)\frac{V(t) - V_{a0}}{V_d - V_{a0}}\left(1 + \frac{1}{Q_{max}}\frac{dV}{dt}(t)\right), \tag{18}$$

where $P_{iso}(t)$ is the active pressure at isovolumic contractions at diastolic volume $V(t) = V_d$, $Q_{max}$ is the maximum blood flow, $V_{p0}$ is the unloaded LV volume, $V_{a0}$ is the "$V_0$" of the time-varying elastance representation of LV, $k_p$ is an elastic modulus.

Let us first consider the approximation of ME models connected in series, and let $P(t)$ be the LV pressure during a single beat of the model. We can then introduce a scaling factor for $\mu(\xi)$ with $\xi \in [0,1]$, which yields

$$P(t) = \mu(\xi)\left(k_p(\bar{V}(t,\xi) - V_{p0}) + P_{iso}(t)\frac{\bar{V}(t,\xi) - V_{a0}}{V_d - V_{a0}}\left(1 + \frac{1}{Q_{max}}\frac{\partial \bar{V}}{\partial t}(t,\xi)\right)\right), \tag{19}$$

where $\bar{V}(\xi)$ are the volumes contributed by the subcomponents of the ME model, with $\xi$ being the discretization variable. Let the global volume for the ME model be $V(t) = \int \bar{V}(t,\xi)d\xi$. We can then represent the elemental volumes as $\bar{V}(t,\xi) = V(t) + \Delta(t,\xi)$, where $\int \Delta(t,\xi)d\xi = 0$. To evaluate the error of approximation, $P(t)$ can then be expressed as

$$P(t) = \mu(\xi^*)\left(k_p(V(t) - V_{p0}) + P_{iso}(t)\frac{V(t) - V_{a0}}{V_d - V_{a0}}\left(1 + \frac{1}{Q_{max}}\frac{dV}{dt}(t)\right)\right) + \epsilon \tag{20}$$

for some $\xi^* \in [0,1]$ with relatively small $\epsilon$. By dividing both sides of (19) by $\mu(\xi)$, integrating with respect to $\xi$, and then by applying the mean value theorem $\frac{1}{\mu(\xi^*)} = \int \frac{1}{\mu(\xi)}d\xi$, we find that error $\epsilon$ in (20) is equal to

$$\begin{aligned} \epsilon &= \mu(\xi^*)P_{iso}(t)\int\left[\frac{\Delta(t,\xi)}{V_d - V_{a0}}\right]\left[\frac{1}{Q_{max}}\frac{d\Delta}{dt}(t,\xi)\right]d\xi, \\ |\epsilon| &\le \mu(\xi^*)P_{iso}(t)\left[\frac{\max_{t,\xi}|\Delta(t,\xi)|}{V_d - V_{ao}}\right]\left[\frac{1}{Q_{max}}\max_{t,\xi}\left|\frac{\partial \Delta}{\partial t}(t,\xi)\right|\right]. \end{aligned} \tag{21}$$

Equation (21) demonstrates that the error of the approximation with respect to $P_{iso}$ is bounded by two terms. The first term of $\Delta$ is $\ll 1$ when heterogeneity is small, as $\max_{t,\xi}|\Delta(t,\xi)|$ is negligible compared to $V_d - V_{a0}$. The second term of $\Delta$ is instead small at large $Q_{max}$ (or high $V_{max}$ of the myofilament model). The latter was illustrated in our simulations as at control values of $V_{max}$ the error is very small but increases with lower $V_{max}$. While we do not provide details for the analysis of $\max_{t,\xi}|\Delta(t,\xi)|$ and $\max_{t,\xi}\left|\frac{d\Delta}{dt}(t,\xi)\right|$, we here note that they depend on te boundaries of $\mu(\xi)$, on the ratio between $k_p$ and the maximum of $P_{iso}$, and on the derivative of $P_{iso}$. For ME models connected in parallel, a similar analysis yields

$$P(t) = k_p(V(t) - V_{p0}) + \int\left[P_{iso}(t)\left(\mu(\xi)\frac{V(t) - V_{p0}}{V_d - V_{a0}} + \frac{V_{p0} - V_{a0}}{V_d - V_{a0}}\right)\left(1 + \mu(\xi)\frac{1}{Q_{max}}\frac{dV}{dt}(t)\right)\right]d\xi. \tag{22}$$

Introducing $\bar{\mu} = \int \mu(\xi)d\xi$, we obtain

$$P(t) = k_p(V(t) - V_{p0}) + P_{iso}(t)\left(\bar{\mu}\frac{V(t) - V_{p0}}{V_d - V_{a0}} + \frac{V_{p0} - V_{a0}}{V_d - V_{a0}}\right)\left(1 + \bar{\mu}\frac{1}{Q_{max}}\frac{dV}{dt}(t)\right) + \epsilon. \tag{23}$$

The approximation error is therefore proportional to the difference

$$d\mu = \left| \left( \int \mu(\xi) d\xi \right)^2 - \int \mu^2(\xi) d\xi \right|, \tag{24}$$

such that

$$|\epsilon| \leq d\mu \times P_{iso}(t) \frac{V_d - V_{p0}}{V_d - V_{a0}} \left[ \frac{1}{Q_{max}} \max_t \left| \frac{dV}{dt}(t) \right| \right]. \tag{25}$$

For example, for the specific $\mu(\xi) = a + b \times \xi$, where $a$ and $b$ are constants, $d\mu = \frac{b^2}{12}$. We can see that the error of approximation again depends mostly on $V_{max}$.

Remarks:

- The approximation error would vanish for elastance-resistance models with constant resistance, e.g., as for the type 2 models in [15]. However, our LO model are more similar in structures to elastance-resistance model with non-constant resistance.
- For simplicity, our analysis considers only linear passive properties and, therefore, additional mismatches might be attributed to nonlinear elasticity, which is instead accounted for in the empirical result section of the manuscript (i.e., see Table 4 in the main text). Discrepancies are expected to increase when nonlinearity is more pronounced.
